# Spontaneous movements and their impact on neural activity fluctuate with latent engagement states

**DOI:** 10.1101/2023.06.26.546404

**Authors:** Chaoqun Yin, Maxwell D. Melin, Gabriel Rojas-Bowe, Xiaonan Richard Sun, João Couto, Steven Gluf, Alex Kostiuk, Simon Musall, Anne K. Churchland

## Abstract

Existing work demonstrates that animals alternate between engaged and disengaged states during perceptual decision-making. To understand the neural signature of these states, we performed cortex-wide measurements of neural activity in mice making auditory decisions. The trial-averaged magnitude of neural activity was similar in the two states. However, the trial-to-trial variance in neural activity was higher during disengagement. To understand this increased variance, we trained separate linear encoding models on neural data from each state. The models demonstrated that although task variables and task-aligned movements impacted neural activity similarly during the two states, movements that are independent of task events explained more variance during disengagement. Behavioral analyses uncovered that during disengagement, movements become uncoupled to task events. Taken together, these results argue that the neural signature of disengagement, though obscured in trial-averaged neural activity, is evident in trial-to-trial variability driven by changing patterns of spontaneous movements.

## Introduction

Latent neurobehavioral states shape natural and trained behaviors in many species^1–4^. Behavioral state may be modulated by a variety of factors, including strategy, arousal, engagement, satiety, task expertise, circadian rhythms, fear, and others^5–8^. Estimating behavioral states has been particularly challenging in animal models due to the latent nature of these states. Recent applications of state-space modeling to behavioral and neural data have granted insights into states that govern decision-making and neural dynamics^3,9–13^. One way to determine an animal’s latent state at a particular moment is to deploy a hidden markov model (HMM) that can assign, for instance, each trial in an experimental session to a specified state. A novel variant of this approach couples the HMM with Bernoulli generalized linear model observations (GLM-HMM)^2,14,15^. This makes it possible to identify latent state fluctuations that govern decision-making dynamics, such as sensitivity to the incoming stimulus and response bias. These models have provided fundamental insights into rodent behavior, revealing that experts performing perceptual decision-making tasks often fluctuate in their engagement levels within single sessions^2,14^. However, it is unknown how these states modulate neural activity during perceptual decision-making.

Previous studies have reported other behavioral indicators of latent states, such as pupil diameter and locomotion^16–20^. Since the inputs to the GLM-HMM include information about the animal’s decisions, it is not entirely surprising that states defined by the model differ in terms of animal performance. It is therefore unclear what other behavioral signatures accompany these model-defined states.

To answer these questions, we used the GLM-HMM to define states in head-fixed mice making decisions about auditory stimuli. We compared the neural activity in these states using widefield calcium imaging. The trial-averaged neural activity was similar in the two states. However, the single trial activity was substantially more variable when mice were in the disengaged state. Using a linear encoding model, we found that we could better explain fluctuations in neural activity during the disengaged state. This increase in explained variance was associated with movements that were not time locked to task events. This difference in neural encoding points to a new hypothesis about state-dependent differences: the alignment of certain movements to task events may be disrupted during disengagement. We made a new behavioral model to quantify the task-aligned and task-independent components of movement. This model revealed a robust negative correlation between task engagement and the magnitude of task-independent movements. Taken together, these observations suggest that task-independent movements not only modulate neural activity in a state-dependent fashion, they are also an indicator of an animal’s latent state of engagement. Importantly, animals make frequent movements when highly engaged in the task; it is not the magnitude of movements that is informative, but instead, their timing and trajectories.

## Results

### Mice occupy engaged and disengaged latent states while performing an auditory perceptual discrimination task

We trained four EMX-Cre-GCaMP6s mice to perform head-fixed auditory discrimination. Mice were head fixed to a behavioral apparatus with speakers on the left and right side of the animal’s head (Fig. 1a, left). Mice initiated trials by grabbing a set of handles in front of them; stimuli were then presented at randomized times after this handle grab. Poisson distributed clicks were played through the left and right speakers^21–23^. After a variable delay period (0 - 1 seconds), two spouts moved in and the mouse reported its decision by licking to the side with more auditory clicks (Fig. 1a, right; Methods, Task 1). The mean rate of left vs right clicks was systematically varied to alter the difficulty of trials. Once mice were trained on all stimulus difficulties, we computed the psychometric function for individual mice (Fig. 1b, light gray lines) and for all mice (Fig. 1b, black line).

**Figure 1:**
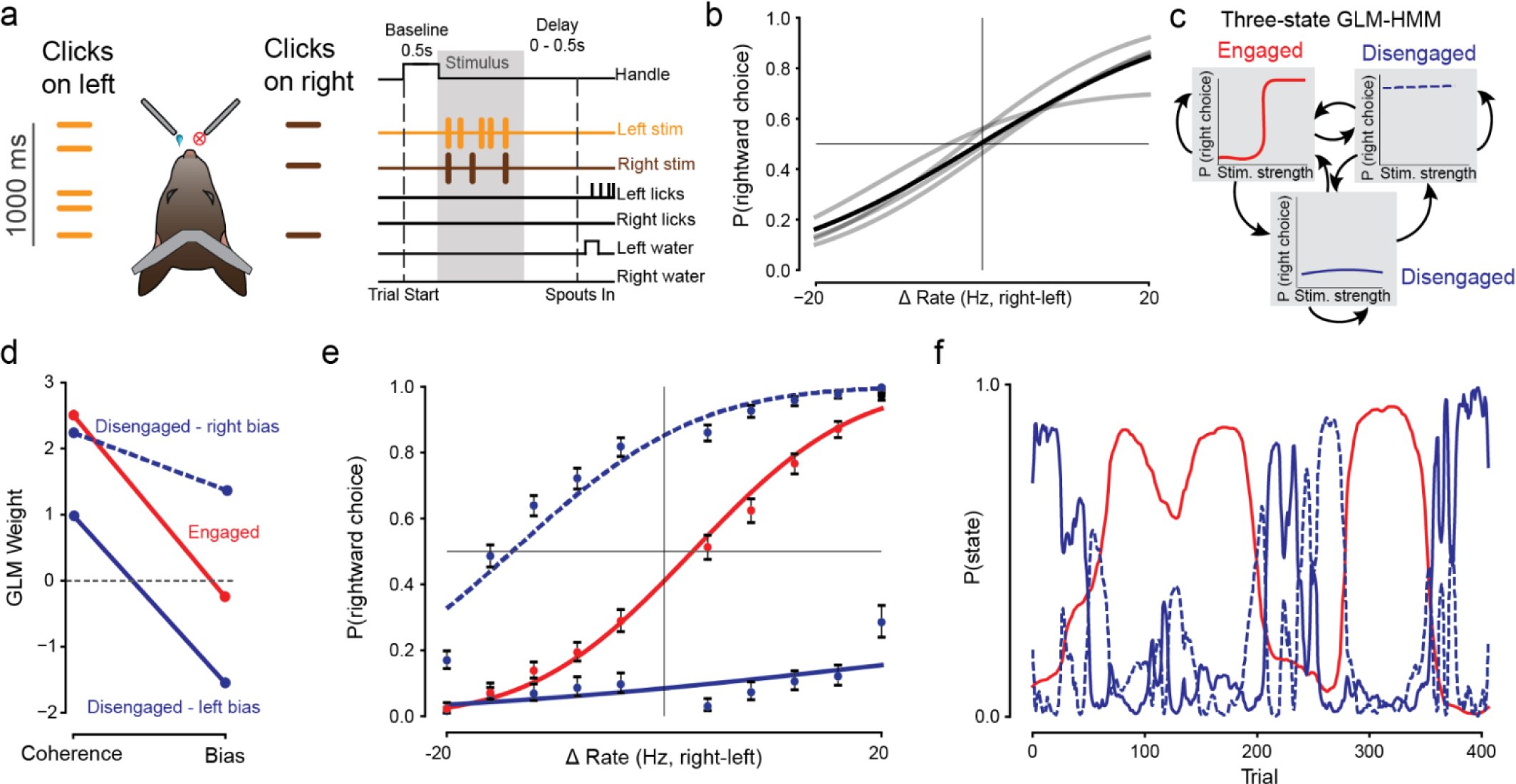
A GLM-HMM identifies discrete behavioral states with distinct psychometric functions. **a)** Head-fixed mice perform a spatial auditory discrimination task. Mice are instructed to lick to the high-rate side after a short delay period. **b)** Psychometric curves for 4 mice (gray lines) and average psychometric curve across all mice (black line). **c)** Example schematic of GLM-HMM. Each latent state has a distinct psychometric curve. Black arrows denote transitions between states. **d)** Parameters from the GLM-HMM fit to the four mice shown in (b). GLM weights for each of the 3 latent states are shown, revealing one engaged and two biased states. **e)** Psychometric curves for each latent state identified by the GLM-HMM. Engaged state: bias=0.131, slope=1.73. Disengaged states: bias=0.936 and -0.699, slope=0.617 and 1.49. Error bars indicate Wilson Binomial Confidence Intervals. Same color conventions as in (d). **f)** Recovered state probabilities from an example session, revealing fluctuations in engagement throughout a session. Same color conventions as in (d).

We fit a three-state GLM-HMM (Fig. 1c) to the behavioral data from this task. To ensure sufficient trial counts for the model, we combined data from the four mice shown in Figure 1b. The model identified one engaged state with high stimulus weight, and two disengaged states (right and left biased, Fig. 1d). After fitting the GLM-HMM, we used the forward-backward algorithm, which computes the posterior probabilities of hidden Markov states given a sequence of observations, to compute the probability that a mouse was in a particular state for a given trial. We then assigned trials to three groups, identified by their most likely state, and computed the psychometric function for each state (Fig. 1e, Methods). As expected, the engaged state showed the lowest bias and the highest slope. Disengaged states were marked by noticeable rightward or leftward bias and a smaller slope value. Fluctuating state probabilities for an example session are shown in Fig. 1f, demonstrating state transitions that span tens to hundreds of trials. For subsequent analyses, we grouped trials into either engaged or disengaged states (where trials from left- and right-biased states were combined).

### Trial-averaged cortical activity is similar across states, but variability across trials increases during disengagement

Armed with a tool to designate trials as originating from engaged or disengaged states, we next tested whether these latent states might modulate neural activity. To do this, we examined widefield imaging signals from transgenic mice expressing GCaMP6s in all excitatory neurons^22^ (Methods). Fluorescence was measured through the intact, cleared skull. Fluorescence data were spatially aligned to the Allen Mouse Common Coordinate Framework v3 (CCF)^24^. Data were temporally aligned to task epochs (baseline, handle-grab, stimulus, delay, response) to compute individual trial responses and trial-averaged responses.

To understand the impact of engagement on neural activity, we first examined the neural responses of trials assigned to the engaged or disengaged state. Trials were assigned to disengaged or engaged states by applying a probability threshold of 0.8. Thus, if the probability of a trial being from a particular state was greater than 0.8, then that trial was assigned to the corresponding state (Fig. 2a). Within sessions, we sampled equal numbers of trials from each state. Sessions with fewer than 25 trials in either state were excluded from analysis. In the somatosensory cortex, upper limb area (Fig. 2b), we found that in both states, the trial-averaged neural activity fluctuated as the animals initiated trials, formed decisions, and obtained rewards (Fig. 2b thick traces). Considerable single-trial variability was also apparent (Fig. 2b, thin traces). For this and subsequent analyses, we matched the ratio of correct to incorrect outcomes for the trials that were studied in each state (Fig. 2c). This ensured that any state-dependent differences we observed in neural activity were not due to a mismatch in reward-related activity between the two states. Trials in the disengaged state were often either right- or left-biased and pooling these ensured that both left- and right-choice trials were included in the analysis. An imbalance between left- and right-trials was apparent in both states.

**Figure 2:**
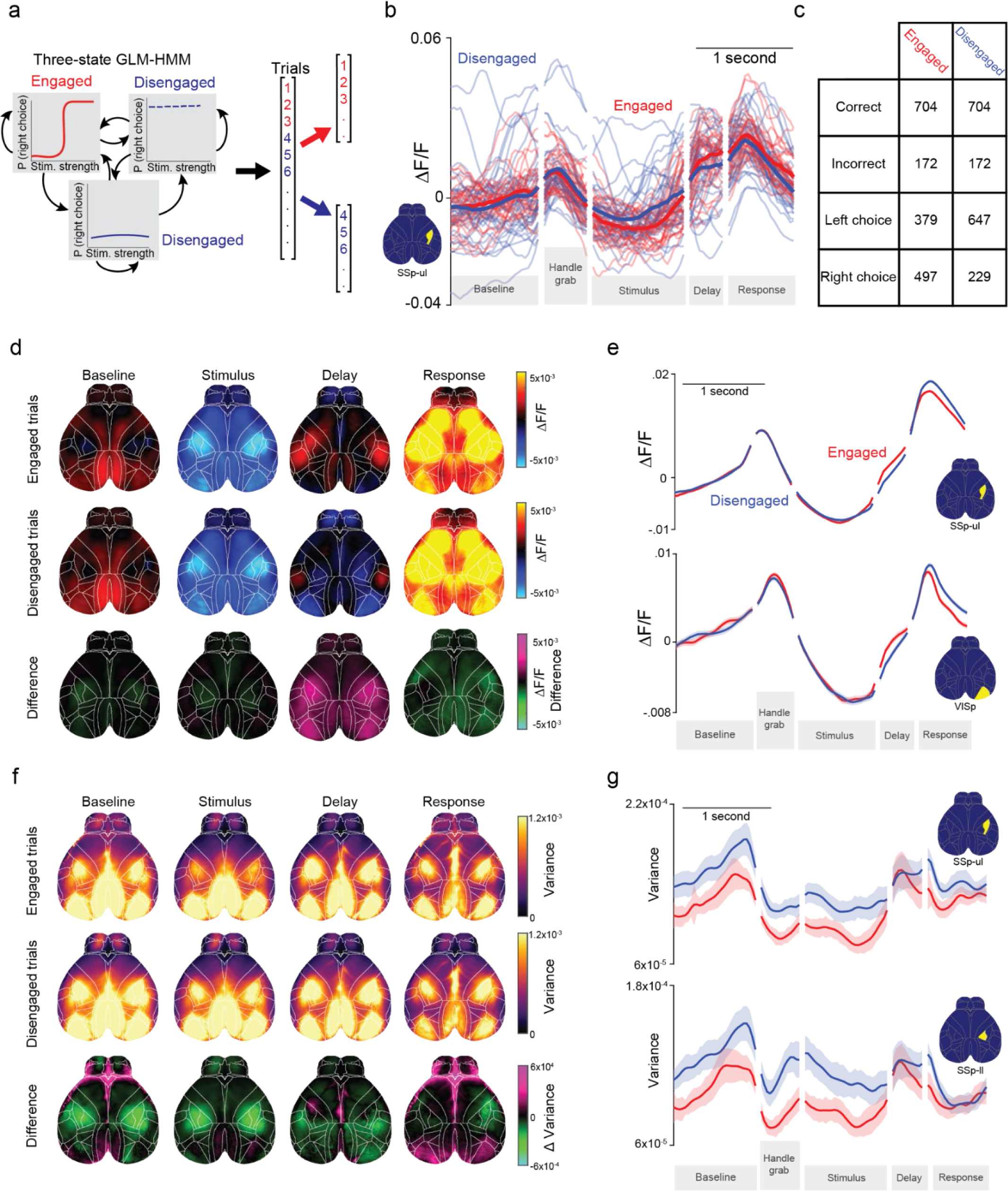
Trial-average responses measured with widefield imaging are similar, but single trial variability is altered during engagement vs disengagement. **a)** Estimated states from the GLM-HMM were used to group trials into engaged and disengaged states. **b)** Calcium activity from somatosensory cortex-upper limb area (SSp-ul) aligned to task events for an example session. Red lines denote engaged trials, blue lines denote disengaged trials. Thin lines: activity on individual trials. Thick lines: mean trial-averaged neural activity. **c)** Trial numbers for the analyses shown in d-g. Rewarded/unrewarded trials are counterbalanced to ensure state-dependent differences are not driven by reward. **d)** Heatmaps of average activity during each trial epoch, averaged over 19 sessions from 4 mice. Top row: engaged trials. Middle row: disengaged trials. Bottom row: difference (engaged activity minus disengaged activity). Purple indicates more activity during the engaged state, green indicates more activity during the disengaged state. **e)** Average activity over time for two example regions. Error: +/- SEM across sessions. **f)** Heatmaps of average variance across trials. Higher values denote more trial-to-trial differences in neural activity. Row conventions are the same as in (e). **g)** Across-trial-variance plotted for two example regions. Error: +/- SEM across sessions.

We then compared mean neural activity across states, not for only a single area (Fig. 2b) but for the entire dorsal cortex (Fig. 2d). We considered 4 time periods in the trial: baseline, stimulus, delay and response (see gray boxes in Fig. 2b). The mean activity for engaged (Fig. 2d, top) and disengaged (Fig. 2d, middle) trials were similar at most timepoints, with only transient differences between the two (Fig. 2d, bottom). A closer look at the temporal dynamics of mean activity in somatosensory and visual cortices showed that trial-averaged neural activity on engaged trials was slightly elevated during the delay period (Fig. 2e, red trace above blue trace during “delay”).

Having observed only modest state-dependent differences in the trial-averaged response (Fig. 2d,e), we then computed the cross-trial variance. This analysis is important because features that modulate neural activity at different moments in each trial will be obscured when many trials are averaged together to generate the mean response. Interestingly, we found that the cross-trial variance on disengaged trials was reliably higher (Fig. 2f). This means that single trial activity from the disengaged state is less stereotyped than in the engaged state, a feature which can be seen when examining single-trial responses (Fig. 2b, compare thin blue/red traces). This state-dependent difference in trial-to-trial variability was evident across multiple task epochs, including baseline, initiation, and stimulus periods. These differences were most evident in primary motor cortex (MOp) and somatosensory cortex (specifically the upper limb area, lower limb area, and barrel cortex, Fig. 2g). Interestingly, we also noticed the opposite effect in posterior visual areas during only the response epoch, where disengaged trials were briefly less variable. Because we counterbalanced correct and incorrect trials within sessions, these findings are not due to a difference in reward rate. These findings were also not due to the state-dependent left and right choice imbalances shown in Figure 2c. When equalizing left and right choice trials within each state (instead of rewarded vs unrewarded ratio) we found that the engaged state was again associated with less trial-to-trial variability (Supplementary Figure 1). Taken together, these results suggest that cortical activity is more stereotyped during engaged, high-performance behavior. In contrast, disengagement is associated with a greater trial-to-trial variability in cortical activity.

### Task variable encoding remains intact, while movement encoding is altered during disengagement

The analyses above demonstrate that cortical activity is more variable during disengagement, but leave unclear what factors contributed to this increased variability. To compare the relationship between movements/task variables and single trial activity during the two states, we used a linear encoding model. We trained separate models on trials that the GLM-HMM assigned to the engaged vs. disengaged states (Fig. 3a). Engaged- and disengaged-state models were matched within sessions to have the same amount of training data. These models included task variables (e.g., the onset time of the stimulus, reward), discrete movements (e.g., whisks, licks) and video data. These models were able to reliably reconstruct single-trial activity across the dorsal cortex (Fig. 3b). To assess overall model performance, we determined the match between the model’s prediction and the actual neural data, using data held out from the training (cross-validated R^2^, Fig. 3c). This cvR^2^ calculation was done separately for each state. Interestingly, this metric was higher for models trained on disengaged trials (Fig. 3c, left, blue line above red line), meaning that the model could account for more single-trial variance in neural activity when animals were disengaged.

**Figure 3.**
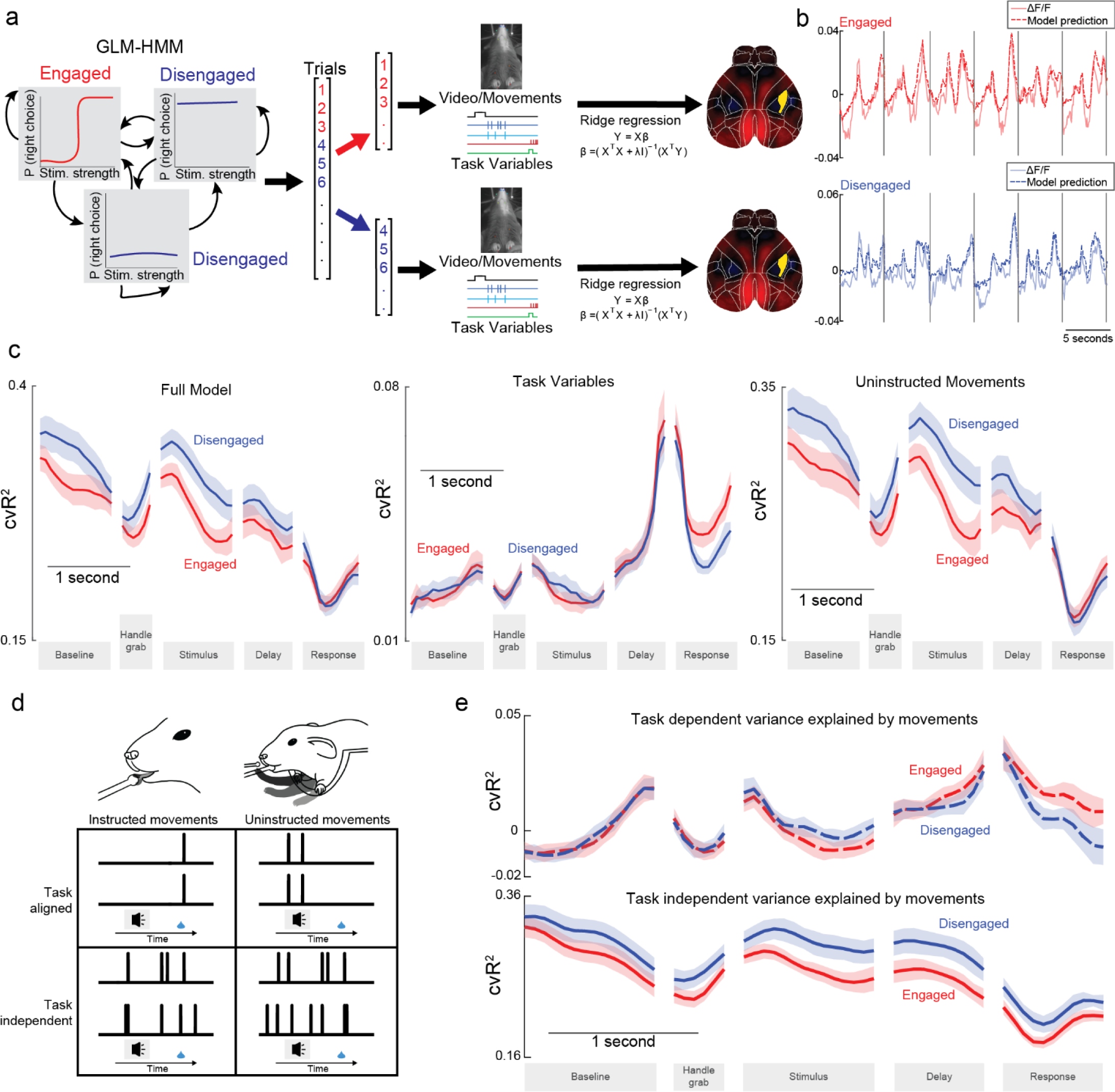
More single-trial variance can be explained in the disengaged state, primarily because of variance due to task-independent, uninstructed movements. **a)** Schematic describing the modeling approach. Trials were identified as engaged or disengaged with a GLM-HMM. Linear models used video, movements, and task variables to predict neural activity. For each session, data from engaged and disengaged trials were fed into separate models. Ridge regression was used to predict single trial neural activity from the input regressors. Cortical heatmaps on the right denote instantaneous ΔF/F to be predicted by the model, where red indicates positive ΔF/F and blue indicates negative ΔF/F. To ensure models were well-fit, sessions with fewer than 50 trials in each state were excluded from analysis. **b)** Example traces from SSp-ul (yellow highlighted region from (a)) and model predictions for models trained on engaged and disengaged trials from one session. Grey vertical lines denote the beginning of a trial. **c)** Total variance explained across trial epochs. Variance explained is averaged across the whole dorsal cortex. Left, variance explained for full model. Middle, variance explained for model containing only task variables. Right, variance explained for model containing only uninstructed movements. Data were combined across sessions with the mean +/- SEM plotted. **d)** Diagram demonstrating how movements can be viewed as instructed vs uninstructed or task aligned vs task independent. **e)** Same as (c), but variance explained is shown for task dependent (top) vs task independent (bottom) movements.

The difference in variance explained could occur if task variables, such as the stimulus or a decision variable, are less reliably encoded during disengaged states, leading to greater variability in activity across trials. This was not the case for the majority of the trial; when we repeated the analysis with a simplified model containing exclusively task variables, we observed no difference in model performance on the engaged vs. disengaged states for most of the trial (Fig. 3c, middle). An alternative explanation is that movements drove the trial-to-trial variability during disengagement. We trained another set of models using only uninstructed movement and found that we were again able to better predict neural activity during disengagement (Fig. 3c, right, blue line above red). This indicates that movements are the primary driver of the previously discussed state-dependent differences.

In this behavioral task, previous work suggests that about 83% of model-explained variance in cortical widefield activity comes from trial-by-trial variability, while around 17% of explainable variance is due to condition-averaged activity (the PETH for each trial condition)^23^. This work also demonstrated that movements can contribute to both trial-by-trial variability and condition-averaged activity: this is because the space of all movements includes task-aligned and task-independent components (Fig. 3d, compare top vs. bottom rows)^23^. We wondered if the observed state-dependent differences were driven primarily by task-aligned movements (i.e. neural variance that can be explained by task variables *or* movements, due to correlation between the two) or task-independent movements (i.e. neural variance that can only be explained by movements, which are uncorrelated to task variables). Using previously described methods^23^, we computed the variance accounted for by these two types of movements: when movements and task variables are correlated, they account for overlapping cortical variance in a linear encoding model. This correlation can be exploited by comparing the explained variance in models with only task variables vs. models with task variables plus movements. The additional variance explained by including movements represents task-independent variance (variance due to movements that cannot be accounted for by task variables). Because movement-driven variance must be either task-aligned or task-independent, the task-aligned variance is then computed by subtracting task-independent variance from the total variance explained by movements.

Movements that were aligned to task events explained equal amounts of cortical variance in engaged and disengaged states (Fig. 3e, top row, compare red/blue dashed traces, see Methods). These include instructed and uninstructed movements that are aligned in time to stimulus events, and they accounted for a small amount of variance that was similar during the two states. However, when we compared model performance on movements that were independently timed relative to task events, we found a consistent difference between the two states (Fig. 3e, bottom row, blue line above red line).

This analysis suggests that the impact of latent state on neural activity is related to the timing of movements. However, the analysis falls short of uncovering what aspects of the animal’s movements are changing: our model comparison can only show that there are changes in the magnitude of explainable cortical variance, but leaves unanswered how the movements were altered during disengagement to drive this change.

### Overall movement magnitude is consistent across states

To better understand movements in the engaged vs. disengaged states, we developed a new analysis, based on the movement of individual body parts. We employed DeepLabCut, a neural-network-based movement tracking software^25^, to track the movements of 27 body parts during the task (Fig. 4a, Supplementary videos 1 & 2). We used DeepLabCut tracking instead of dimensionality-reduced videos or raw pixel values for two reasons. First, state-dependent differences in dimensionality-reduced video are difficult to relate to specific animal movements. Second, using raw pixel values can lead to ambiguities: for instance, a large movement of a small body surface and a small movement of a large body surface can lead to identical frame-to-frame changes in video pixels.

**Figure 4:**
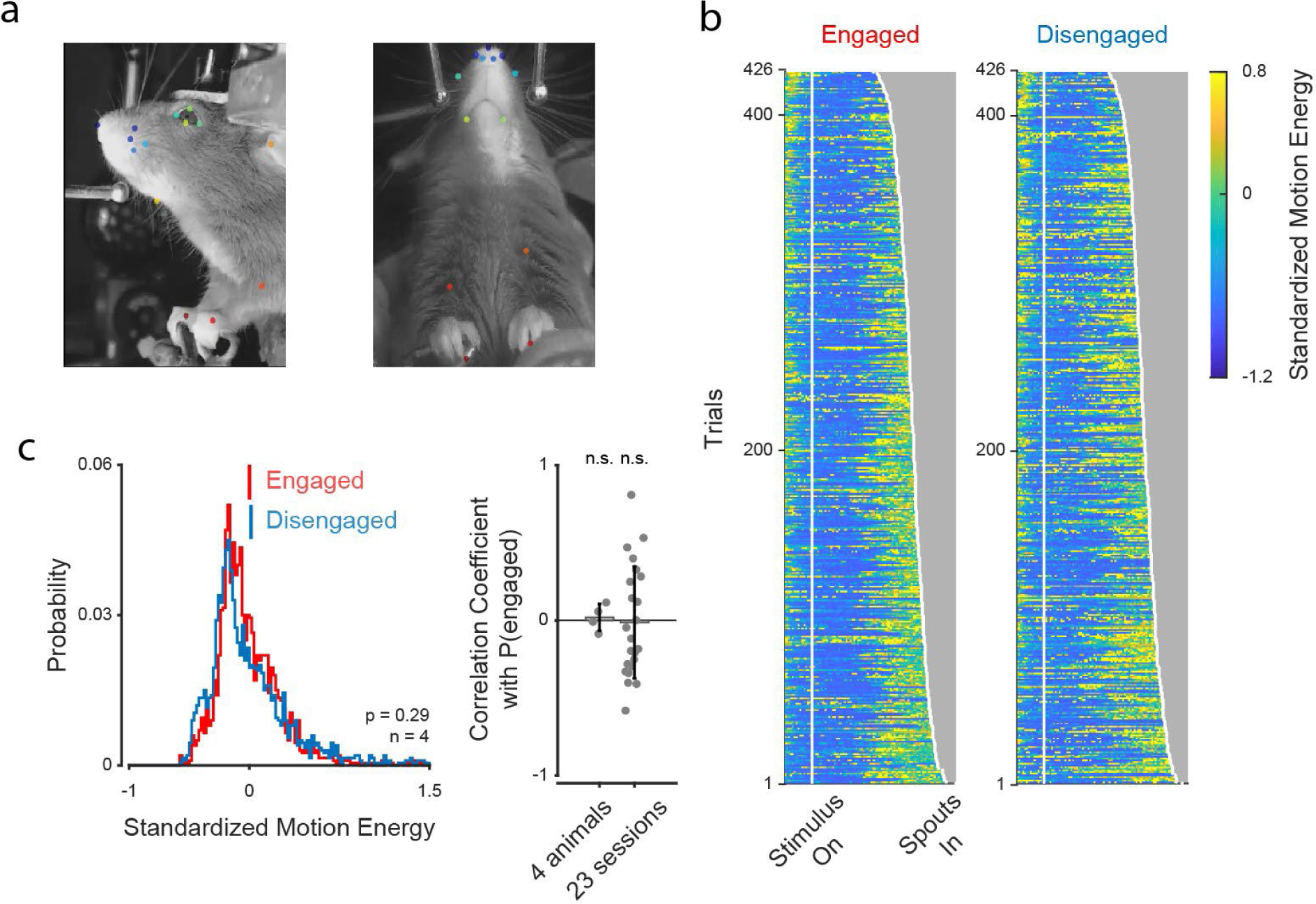
Animals move the same amount in the engaged and disengaged states. **a)** The movements of 27 body parts were tracked with DeepLabCut. Each panel shows an example frame from one of the two cameras used to track the animal’s movements. Colored dots: points tracked by DeepLabCut. **b)** The motion energy of the DeepLabCut labels for an example animal on engaged (left) and disengaged (right) trails. Each row is one trial. Trials are sorted based on the time that the water spouts moved in and were available for the mouse to report a choice (“spouts in”). **c)** The motion energy is similar across engaged and disengaged states. Left panel: the distribution of motion energy (averaged over the stimulus and delay windows for each trial). The red/blue bars denote the mean motion energy of engaged/disengaged states, respectively. (Standardized motion energy of engaged trials = 0.0022 (mean) ± 0.3075 (standard deviation), standardized motion energy of disengaged trials = 0.0143 ± 0.3965, unpaired t-test, p = 0.2889. 3836 trials from 4 animals were included.) Right panel: the correlation coefficient between motion energy and P(engaged) was calculated separately for individual mice (left) and for individual sessions (right). Bars denote the averaged coefficients across mice and sessions. No correlation exists between motion energy and the probability of being in the engaged state (P(engaged)). (Correlation coefficients of 4 mice = 0.0193 ± 0.0870, these values are not significantly different from zero, one-sample t-test, p = 0.6870; Correlation coefficients of 23 individual sessions = -0.0132 ± 0.3599, these values are not significantly different from zero, one-sample t-test, p = 0.8622)

To test whether mice move more during disengagement, we quantified the motion energy (cumulative position change across frames) of labeled body parts in the stimulus and delay epochs. To prevent the total motion energy from being dominated by body parts that naturally have longer moving distances, such as the forelimbs, we standardized (z-scored) the motion energy of different body parts separately before averaging them together. In most trials, standardized motion energy tended to increase gradually from the beginning of the trial to the moment when the water spouts came in and the mouse reported its choice (Fig. 4b). Interestingly, the motion energy was similar during engagement and disengagement (Fig. 4c). This argues that the increase in explained variance during disengagement (Fig. 3c,e) did not arise because disengaged animals move more.

### Stereotyped movement patterns are disrupted during disengagement

Next, we tested whether the timing of the movements changed during disengagement. The timing of movements could change if, for instance, movements became desynchronized to task events during disengagement. We have previously observed that in trained experts (but not novices), animals make well-timed whisks when the stimulus begins and when the reward is delivered^22,23^. To quantify the extent to which movements are aligned in time to task events, we built a model that used task variables (e.g. stimulus strength, upcoming choice) to predict the movement of the DeepLabCut labels (Methods). Our model uncovered, unsurprisingly, that movement patterns differed depending on the task variables (e.g, in advance of left vs. right choices, Fig. 5a; the thick gray trace indicating trial-averaged movement trajectory differs for trials that end in a left vs. right choice; example R^2^ values shown in Supplementary figure 2). However, some movements could not be explained by task events; for instance, a forepaw flexion that occurs randomly from trial-to-trial (Fig. 5a, thin traces) cannot be accounted for by the animal’s choice or stimulus strength. We refer to these as “Task Independent Movements’’ (TIM). We obtained one value of TIM for each trial, capturing the extent to which, on that trial, movements were misaligned to task events (Methods).

**Figure 5:**
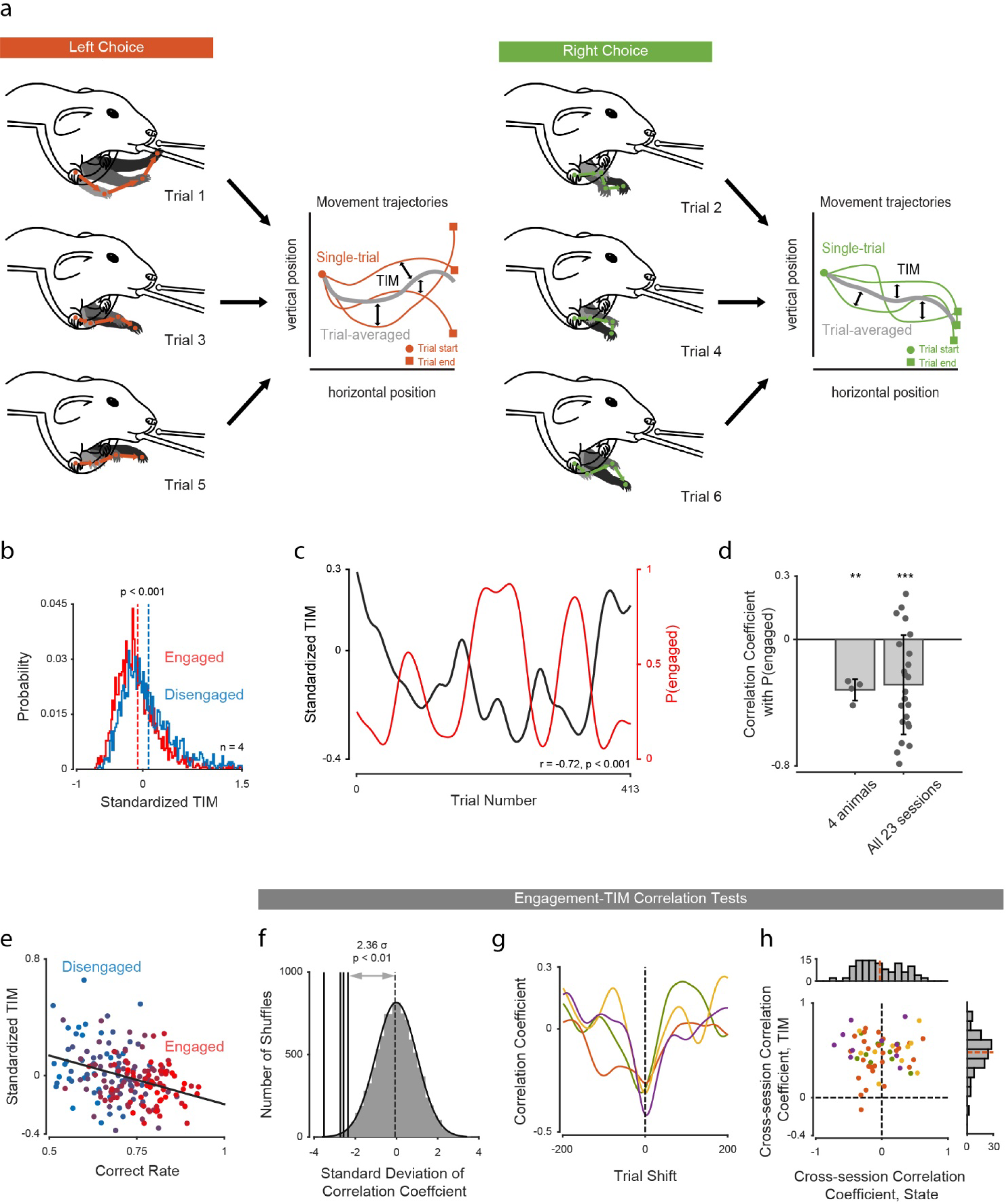
A novel behavioral metric demonstrates a strong relationship between the timing of movements and the animal’s latent state. **a)** The movement of one front paw has a stereotyped trajectory that differs for left choices (left images) and right choices (right images). Task-independent movement (TIM) measures the mismatch between model-predicted stereotyped trajectories and actual single-trial movements. TIM is low when an animal’s spontaneous movements are more stereotypical from trial-to-trial. **b)** The distributions of trial-averaged values of TIM (calculated during the stimulus and delay epochs) in the engaged/disengaged states. The red/blue dashed lines denote the means of the two distributions respectively. TIM is significantly higher in the disengaged state. (Standardized TIM of engaged trials = -0.0755 (mean) ± 0.3387 (standard deviation), standardized TIM of disengaged trials = 0.0860 ± 0.4322, p of unpaired t-test = 3.4994e-37, 3836 trials from 4 animals were included.) **c)** Smoothed TIM and P(engaged) in an example session (r = -0.7163, p = 3.3286e-66). **d)** Correlation coefficients of TIM and P(engaged) were calculated separately for individual mice or sessions after TIM and P(engaged) were smoothed with a 50-trial-long gaussian filter. Bars denote the averaged coefficients across mice and sessions. TIM and P(engaged) were negatively correlated at both single-animal and single-session levels. (Correlation coefficients of 4 mice = -0.3196 (mean) ± 0.0679 (standard deviation), Correlations were significantly less than zero, one-sample t-test, p = 0.0025; Correlation coefficients of 23 sessions = -0.2869 ± 0.3146, one-sample t-test, p = 2.4235e-04) **e)** Scatter plot shows the correct rate and TIM calculated in an independent 50 trial window. The color grade indicates the P(engaged) value of trials. TIM is negatively correlated with both engagement and task performance. (TIM-correct rate correlation: r = -0.3315, p = 2.6445e-06; TIM-P(engaged) correlation: r = -0.2799, p = 8.4443e-05). **f)** The distances between actual correlation coefficients and the Gaussian control distribution (the correlation coefficients of 10,000 shuffled TIM and P(engaged) pairs) confirm that negative TIM-engagement correlations are significant. The actual correlation coefficients of 4 mice are -2.3572, - 2.6434, -2.5707, and -3.5154 standard deviations away from the control distribution respectively. **g)** Trial shift test shows that there is a near-zero time lag in the negative TIM-engagement correlation. Each curve indicates, for a single mouse, the correlation between TIM and p(engagement) when they were offset in time by the value indicated on the horizontal axis. The negative correlation is at its strongest level when the offset was near zero. **h)** Each point denotes the correlation coefficient of a pair of sessions from the same mouse. 4 colors denote 4 individual mice (same color conventions as in (g)). Marginal histograms show the distribution of cross-session correlation of state (top) or TIM (right). Red dashed lines: mean of each distribution. The cross-session correlation of TIM is non-zero (TIM cross-session correlation coefficients = 0.4676 ± 0.1957, one-sample t-test, p = 1.4923e-49, 23 sessions from 4 animals). The cross-session correlation of P(engaged) is not significantly different from 0 (coefficients = -0.0333 ± 0.3216, one-sample t-test, p = 0.2673, 23 sessions from 4 animals).

We then tested whether TIM depended on the engagement states of animals. We first observed that TIM was higher on disengaged trials (Fig. 5b). Moreover, when we compared fluctuations in TIM with those of engagement, we found them to be inversely correlated (Fig. 5c,d). This demonstrates that the periods of low TIM, when movements tend to be better aligned to task events, are associated with engagement on the task. This relationship was present at both overall and the single animal/session levels (Fig. 5d). Accordingly, the spatial distribution of DeepLabCut labels occupies a larger area in high-TIM trials (Supplementary figure 3). This indicates that body label positions are more variable across trials during disengagement. Given that engagement states were modeled based on the animals’ choice, we also tested the relationship between task performance, TIM, and P(engaged). These metrics are related (Fig. 5e and Supplementary figure 4). The engaged state is characterized by both high correct rate and low TIM (Fig. 5e, red points at the right).

To ensure that the negative correlation between TIM and P(engaged) did not occur by chance^26^, we recalculated the correlation coefficient with a version of TIM in which the trial order was randomly shuffled. This step was repeated 10,000 times for each mouse, which generated a Gaussian distribution of “control coefficients”. We thus could calculate the p-value of each correlation based on the distance between the real coefficient and the control distribution. The negative correlations of all 4 animals are significant (Fig. 5f). Moreover, we tested the time lag of TIM-state correlation by shifting TIM over time. The negative correlations of all 4 animals reached their peak around 0, implying that there is a near-zero time lag between TIM and engagement (Fig. 5g).

One possible concern is that the fluctuations of TIM and P(engaged) within each session might have fixed patterns. For instance, TIM might always be higher at the beginning and the end of each session, and P(engaged) might have an opposite pattern. In this case, the correlation between TIM and P(engaged) does not indicate a real association between them. To address this concern, we calculated the cross-session correlations of TIM and P(engaged) respectively after adjusting all sessions to the same length by 1-dimensional interpolation. Namely, if we had 3 sessions from one mouse, we would calculate the TIM correlation of 3 session pairs: session 1 - session 2, session 1 - session 3, and session 2 - session 3. If TIM fluctuation was highly stereotyped within each session, the correlation coefficients of most pairs would be high. We repeated this process for P(engaged). Only TIM showed high cross-session correlation (averaged r = 0.47, 4 animals). The cross-session correlation of P(engaged) was minimal (averaged r = -0.03, 4 animals) (Fig. 5h, marginal histogram at top). This result argues against the possibility that TIM-state correlation emerged from the fixed patterns of both TIM and P(engaged).

A second possible concern is that the linear regression models used to predict the body part positions included outcome-related regressors. The negative TIM-state correlation would then merely reflect the fluctuation of model-fitting quality with specific outcome-related regressors (e.g. rewarded vs unrewarded). To address this concern, we recalculated TIM after excluding all outcome-related regressors and repeated the correlation analyses. The correlation structure between task performance, TIM, and P(engaged) was preserved (Supplementary figure 5).

A final concern is that the increase in TIM may not indicate the animal’s engagement in the current trial, but instead simply reflect that the animal experienced an omitted reward and a time-out punishment in the previous trial. In this case, TIM could be higher in the disengaged state simply because the disengaged state had more incorrect previous trials. However, we observed that TIM was higher for disengaged than for engaged trials even when the outcome of the previous trial was matched (Supplementary figure 6).

### TIM is more tightly linked to cognitive state than pupil diameter

Pupil diameter has previously been used to estimate the engagement level of subjects across species. Previous studies have reported positive or inverted-U-shape correlations between animal engagement and pupil diameter^20,27,28^. We tested whether TIM and pupil diameter provided similar estimates of animal engagement. We extracted the mean pupil diameter in the baseline period (the 0.5s window preceding the handle grab) and stimulus epochs from videos (Fig. 6a), then smoothed and downsampled the data with a 50-trial window. The pupil diameter in neither the baseline nor the stimulus epochs correlated with task performance or P(engaged) (Fig. 6b). We further tested the correlation between pupil dilation (the pupil diameter difference between the stimulus and the baseline epochs), task performance, and P(engaged). No significant correlations were observed (Supplementary figure 7).

**Figure 6:**
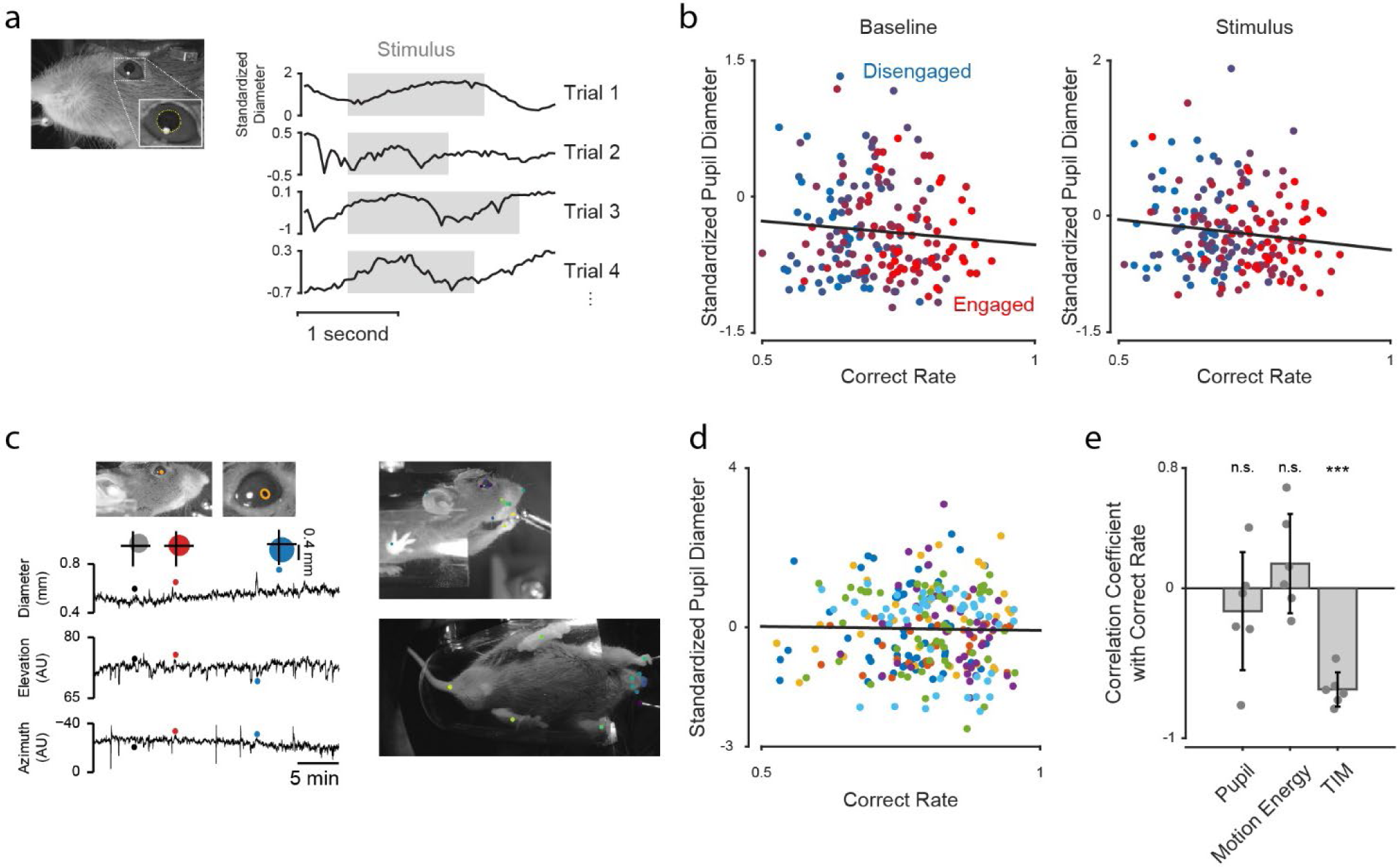
There is no significant association between latent state and pupil diameter. **a)** Pupil diameter was measured in head-fixed mice performing an auditory decision-making task. Left and inset: pupil diameter was measured based on the lateral videos with customized code^22^. Right: the standardized pupil diameter of 4 example trials. Pupil diameters during the baseline and stimulus epochs were used in subsequent analyses. Gray boxes indicate the stimulus epoch of each trial. **b)** Pupil diameter and correct rate were smoothed with a 50-trial long Gaussian filter and downsampled every 50 trials. Each point denotes one trial. The color grade indicates the P(engaged) value of trials. No correlation exists between pupil diameter and task performance/engagement (Baseline period: r = - 0.0881, p = 0.2257; Stimulus period: r = -0.1325, p = 0.0663. n = 4). **c)** Top-left: the pupil diameter was extracted from the lateral videos of head-fixed mice performing a visual task. Pupil diameter was estimated using customized software^29^. Bottom-left: Filtered pupil diameter (area shown as circles) and eye position (crosshairs represent the center of the eye). Right: 30 body parts in the new dataset were tracked with DeepLabCut for motion energy and TIM calculation. The colored dots show the DeepLabCut labels on two example frames from the videos. **d)** Relationship between correct rate and standardized pupil diameter for 6 mice (colors) that comprise the dataset described in (c). Correct rate and pupil diameter were smoothed with a 50-trial long Gaussian filter and downsampled every 50 trials to generate the points shown. Pupil diameter and correct rate were not significantly correlated (r = -0.0505, p = 0.3862). **e)** The correlation coefficients of task performance and pupil diameter/standardized motion energy/standardized TIM are plotted for each animal. Bars indicate the averaged correlation coefficients across animals. Only TIM shows significant correlation with task performance. (Pupil-correct rate correlation coefficients = -0.1534 (mean) ± 0.3926 (standard deviation), one-sample t-test, p = 0.3825; motion energy-correct rate correlation coefficients = 0.1640 ± 0.3301, one-sample t-test, p = 0.2781; TIM-correct rate correlation coefficients = -0.6743 ± 0.1147, one-sample t-test, p = 2.9174e-05, n = 6)

Because relationships between pupil diameter and measures of arousal have previously been reported^11,30^, we were surprised to find no relationship here. To ensure that the absent relationship was not limited to a single dataset, we examined a second dataset (Methods, Task 2). Again, we tracked the movement trajectories of 30 body parts with DeepLabCut and extracted the pupil diameter in the 0.5s long window before visual stimuli were displayed (Fig. 6c). We avoided using the stimulus window in this dataset because pupil diameter may fluctuate independently of task engagement when visual stimuli are presented. We again observed no correlation between the pupil diameter and task performance (Fig. 6d). We further tested the TIM-performance correlation and found that, as before (Fig. 5), TIM was inversely correlated with task performance (Fig. 6e).

The significant relationship between TIM and performance we observed suggests that movements might be a more sensitive alternative to pupil diameter as a means to track an animal’s engagement in a task. However, our analyses above were all conducted in head-fixed mice. To test the generality of these observations, we applied TIM-performance analysis to data from freely-moving rats. Rats were trained to distinguish the rate of a 1.0s long multisensory stimulus sequence (see Supplementary figure 8a and Methods, Task 3). We calculated the TIM and motion energy of 19 DeepLabCut-labeled body parts (Supplementary figure 8b). Again, we observed a significant negative correlation between TIM and task performance (Supplementary figure 8c-e).

Combining the results from 3 datasets, we consistently observed that TIM is a stable negative indicator of an animal’s performance on cognitive tasks (Fig. 7). The consistency of this effect across species and contexts argues that spontaneous movements are a reliable indicator of a critical latent variable, the animal’s engagement in a cognitive task.

**Figure 7:**
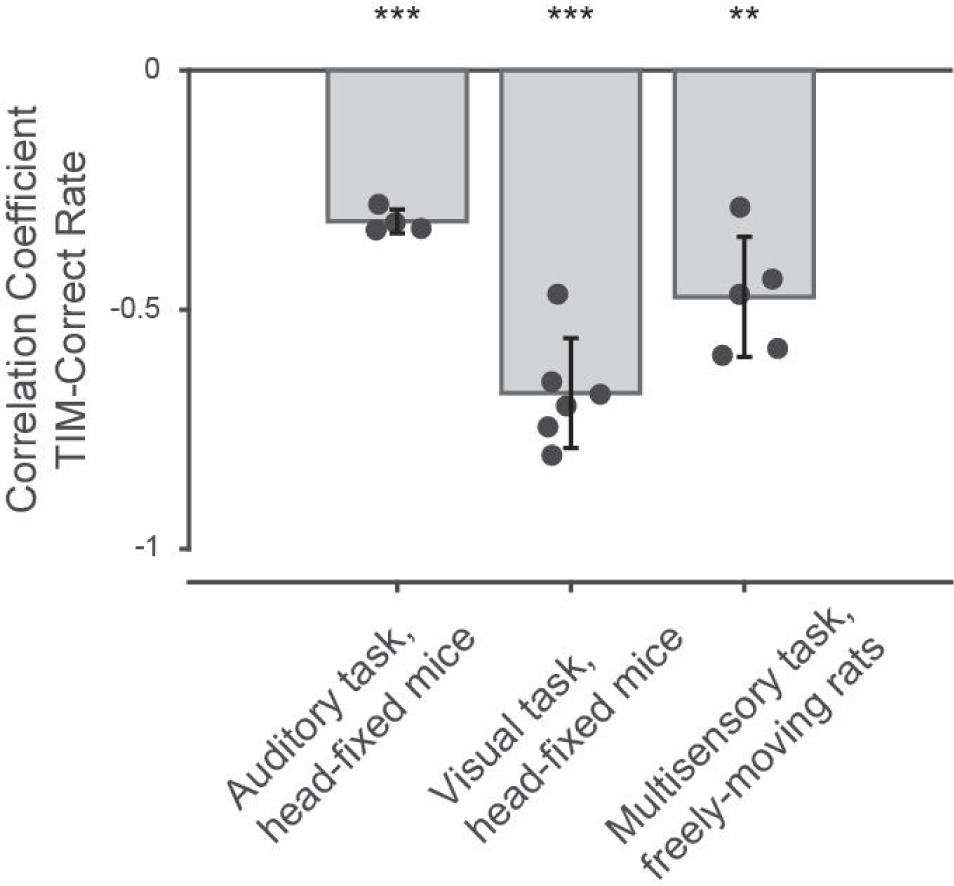
The correlation coefficients between TIM and task performance are plotted for three experimental conditions. Each point denotes the coefficient of one animal. Bars indicate the averaged correlation coefficient across animals. TIM and correct rate are negatively correlated in 3 datasets that span different task paradigms and species. (Correlation coefficients of 4 mice performing the auditory task (task 1) = -0.3156 (mean) ± 0.0247(standard deviation), one-sample t-test, p = 1.3145e-04; Correlation coefficients of 6 mice performing the visual task (task 2) = -0.6743 ± 0.1147, one-sample t-test, p = 2.9174e-05; Correlation coefficients of 5 rats performing the multisensory task (task 3) = -0.4734 ± 0.1254, one-sample t-test, p = 0.0011).

## Discussion

Previous studies have shown that discrete, latent states can govern decision-making dynamics^2^. However, the neural and behavioral correlates that underlie these states remain poorly understood. In this study, we first showed that the trial-averaged activity across the dorsal cortex is largely similar in GLM-HMM-defined states of engagement and disengagement. However, neural activity during the disengaged state exhibited greater variability across trials. Our encoding model analyses further demonstrated that there is a higher proportion of neural activity explained in the disengaged state, driven by the increase in the explanatory power of task-independent movement regressors. These results led us to test whether the variance changes in neural activity were driven by changes in movement dynamics. Multiple studies have shown that even head-fixed animals continuously make spontaneous movements and these movements contribute to a majority of variance in neural activity^23,31,32^. We tracked the movements of body parts with DeepLabCut and quantified the movement differences between latent states. Interestingly, the amount of standardized motion energy was similar in the engaged and disengaged states. However, movements in the engaged state were more temporally aligned to task events. Taken together, these results demonstrate that changes in engagement are associated with altered patterns of spontaneous movements that impact neural activity. This argues that movement patterns can be used as a powerful and versatile indicator of latent states in diverse experimental conditions, and points to a strong link between movements and cognition.

Multiple studies have documented the widespread impact of uninstructed movements on neural activity^23,32–34^, but have left unanswered how these movements impact decision-making. One possibility is that uninstructed movements are *beneficial* to decision-making, especially for decisions that require learning^35^. Indeed, a recent paper shows that mice upregulate spontaneous behavioral syllables when these syllables are paired with dopamine stimulation, suggesting that neuromodulators can allow the brain to sequence spontaneous movements in a structured manner^36^. An alternative possibility is that uninstructed movements *interfere* with decision-making: if movement-encoding and task-encoding subspaces overlap, movements could hinder performance through shared noise or confounding correlations between independent signals^32,37–39^, perhaps along a shared stimulus-movement subspace. Our work paves the way for a third hypothesis: the impact of movements on task execution depends on the nature of the movements. We posit that temporally-aligned movements enhance task-variable encoding along a shared movement-task subspace, and that this facilitates behavior during engagement.

The suppressive effect of engagement on trial-to-trial variability (Fig. 2g) is reminiscent of prior work that likewise observed state-dependent changes in trial-to-trial variability. Those studies showed that trial-to-trial variability drops in a state-dependent way on at least 3 timescales: during the seconds before a learned movement is deployed^40^, during the minutes while attention is deployed to a spatial location^41,42^, and during the weeks-long transition from novice to expert^43^. One possibility is that in those studies, as in ours, the drop in trial-to-trial variability is movement-related. However, a change in movements can’t be the sole explanation because, at least for primate early visual areas, movement-modulation of neural activity is very weak^44,45^. Therefore, the variability drop likely arises, at least in part, from some other source. One hypothesis is that there are modulatory inputs, shared across neurons, that change with the animal’s state^46,47^. An appealing idea is that this same mechanism, a variable shared gain, is present in many species and areas, and the link between this modulatory input and movements is species/areas-specific.

Our data did not show a clear relationship between pupil diameter and decision-making performance. In contrast, previous studies have demonstrated that pupil diameter is related to arousal and task performance^27,28,48–50^ and that this relationship sometimes has an inverted-U shape^8,20,51,52^. The fluctuation of pupil diameter is sensitive to many factors, such as experimental setup (e.g., the light levels in the behavioral rig), behavioral task design (e.g., task difficulty), and physiological factors (e.g., heart rate, locomotion)^53,54^. While we carefully adjusted the lighting in the behavioral setup to allow for a wide range of pupil fluctuations, a possible explanation for why we did not observe the pupil-performance correlation is that our mice were placed on a platform instead of a running wheel. This likely limits the occurrence of high arousal states that are also associated with locomotion^11,20^. Another difference between our study and others is the task structure. While our inter-trial delays were between 1-2s, other studies used inter-trial intervals ranging from 3-7s. Longer inter-trial delays would allow the pupil diameter to recover from reward-induced pupil dilations and for general arousal to decrease before the next trial starts. This is not the case in our task, where mice rapidly make decisions and do not stop working until they are satiated. Thus, one explanation for the inconsistent relationship between pupil diameter and performance observed here is that the pupil diameter in our animals spanned a more restricted range compared to other studies.

More work is needed to better understand the relationship between movements, internal states and neural activity. First, one limitation of this study is that isolating auditory responses within the auditory cortex is difficult with the widefield imaging preparation. Future work that examines state-dependent responses in primary visual cortex to visual stimuli will be able to more definitively determine whether disengagement alters sensory encoding. Second, a study of subcortical structures is needed. Our analysis focused on cortical structures because widefield imaging offers access to many cortical areas simultaneously. However, future work on subcortical structures could be informative. Of particular interest is the posterior nucleus of the thalamus, which has previously shown to be implicated in state-dependent regulation of sensory processing^55,56^. Third, a manipulation experiment could establish a causal link between movements and internal state. Our approach cannot resolve whether TIM drives the animal to distraction or instead reflects a distracted state that was already initiated. Given the difficulty of physically manipulating animals’ movement without abolishing their ability to perform the task, targeted manipulation of state-modulating regions, such as the hypothalamus^57^ or locus coeruleus^51^, could be used to promote an engaged or disengaged state. We could then observe if TIM would reflect such artificial state changes accordingly. Cell-type specific measurements/manipulation are also needed, and perhaps should extend to astrocytes as well as neurons^6,58,59,51,60^.

Previous observations demonstrated that movements have a major impact on neural activity but left unclear how such movements were related to cognition^23,31,32^. Our work uncovers that movements and cognition are intertwined: when animals become engaged in a task, movement patterns change, as does the trial-to-trial variability in neural activity. This points to a close link between movements and cognition and calls for new studies to understand how this link differs across species, areas and behaviors.

## Supplementary figures

**Supplementary figure 1:**
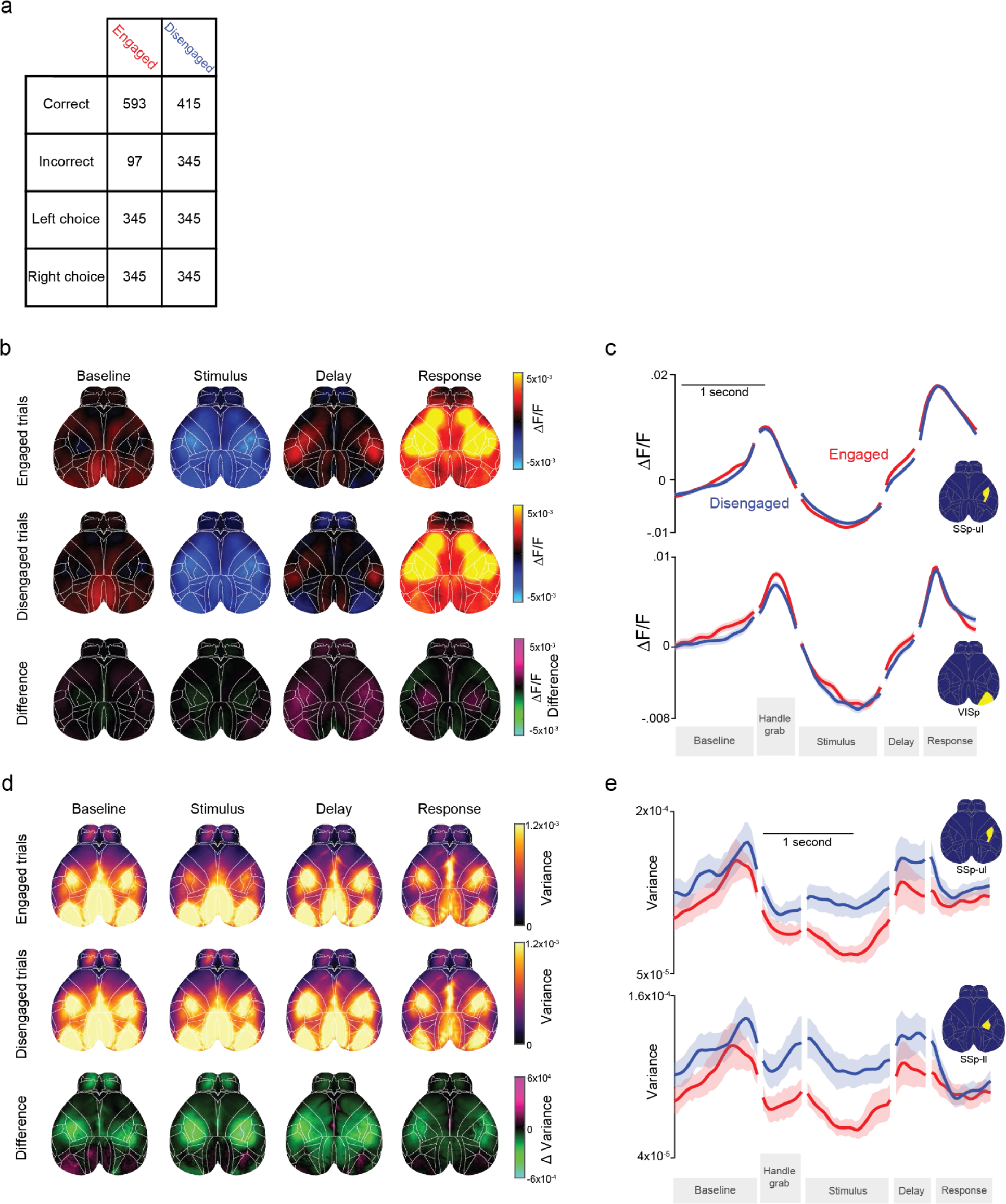
State-dependent differences in cross-trial variance are not due to state-dependent choice differences. **a)** Trial numbers for b-e. Left and right choice trials are equalized to ensure state-dependent differences are not driven by state-dependent choice side preferences. **b)** Heatmaps of average activity during each trial epoch, averaged over 19 sessions from 4 mice. Top row: engaged trials. Middle row: disengaged trials. Bottom row: difference (engaged activity minus disengaged activity). Purple indicates more activity during the engaged state, green indicates more activity during the disengaged state. **c)** Average activity over time for two example regions. Error: +/- SEM across sessions. **d)** Heatmaps of average variance across trials. Higher values denote more trial-to-trial differences in neural activity. Row conventions are the same as in (e). **e)** Across-trial-variance plotted for two example regions. Error: +/- SEM across sessions.

**Supplementary figure 2:**
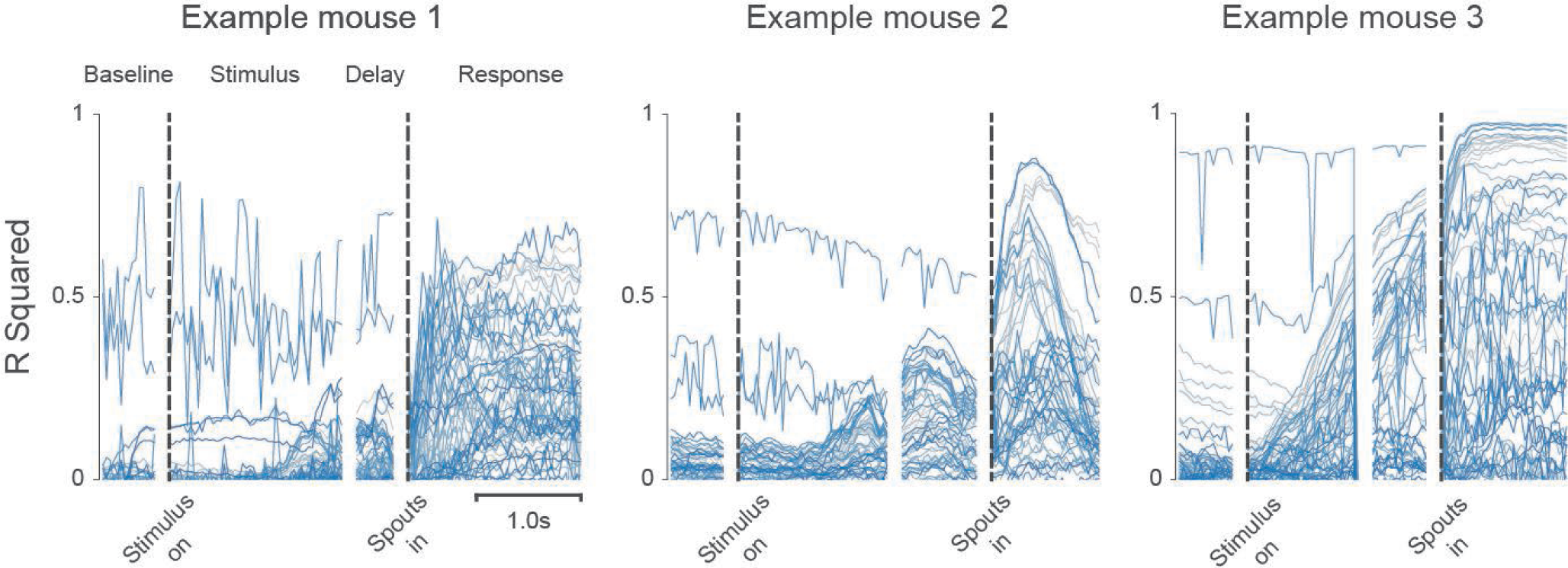
3 panels show the R^2^ values across time of all DeepLabCut labels (x and y are plotted separately) of 3 example animals. Cross-validated linear model captures significant variance in DeepLabCut-tracked movement.

**Supplementary figure 3:**
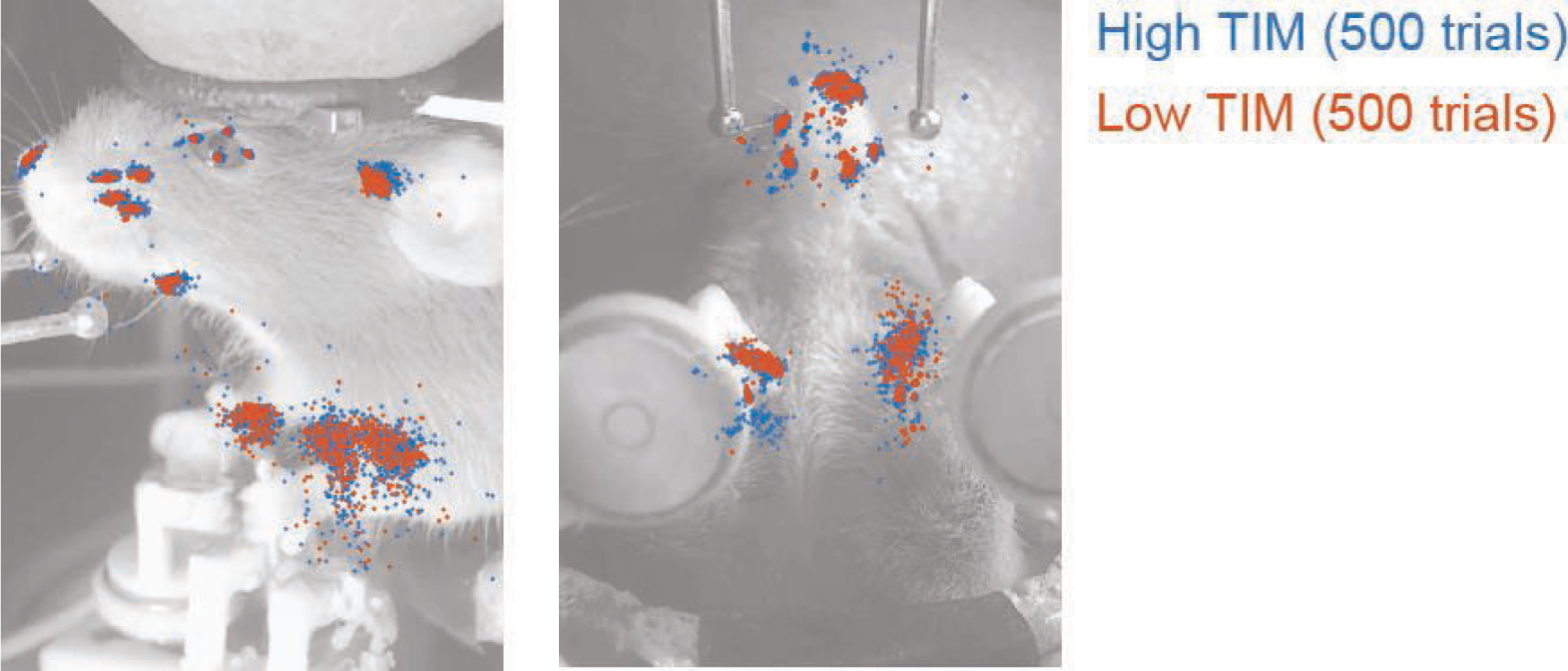
The DeepLabCut label positions of 500 trials with high TIM and 500 trials with low TIM at one example frame (from the stimulus epoch) from one example animal are plotted. The trials with high TIM values distribute in a bigger area.

**Supplementary figure 4:**
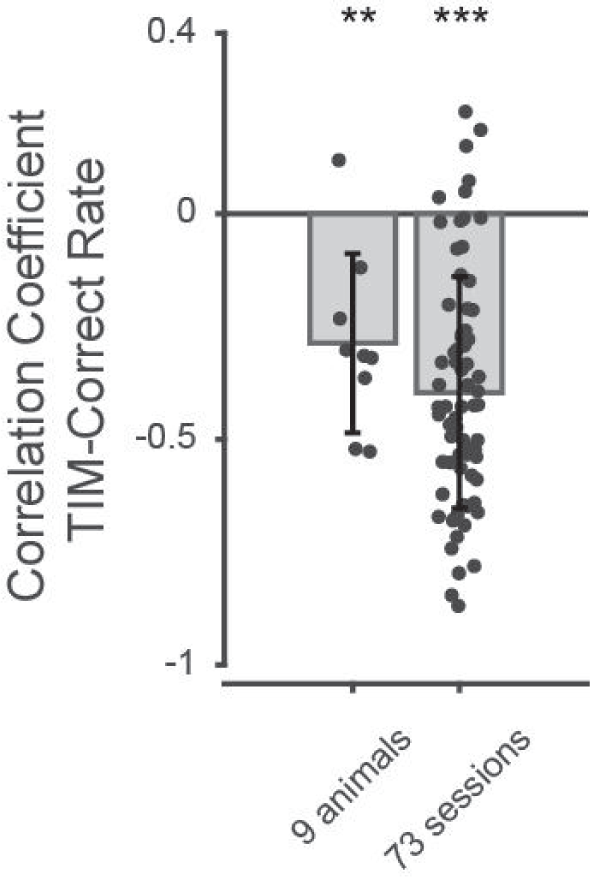
TIM-task performance correlation of 9 additional animals (73 sessions). These animals were used in optogenetic inactivation experiments (inactivation trials would interrupt continuous HMM state) or had cells labeled other than Emx-expression neurons. Thus, these animals were not included in the GLM-HMM analysis. All optogenetic inactivation trials were removed before calculating TIM. As fig. 5d, the correlation coefficients of individual animals (left) and sessions (right) are plotted with dots. The bars indicate the averaged coefficients across animals and sessions. We again observed negative correlation between TIM and correct rate at both individual animal and single session level (Correlation coefficients of 9 mice = -0.2867 (mean) ± 0.1988 (standard deviation), p of one-sample t-test = 0.0025; Correlation coefficients of 73 sessions = -0.3962 ± 0.2566, p of one-sample t-test = 6.8090e-21).

**Supplementary figure 5:**
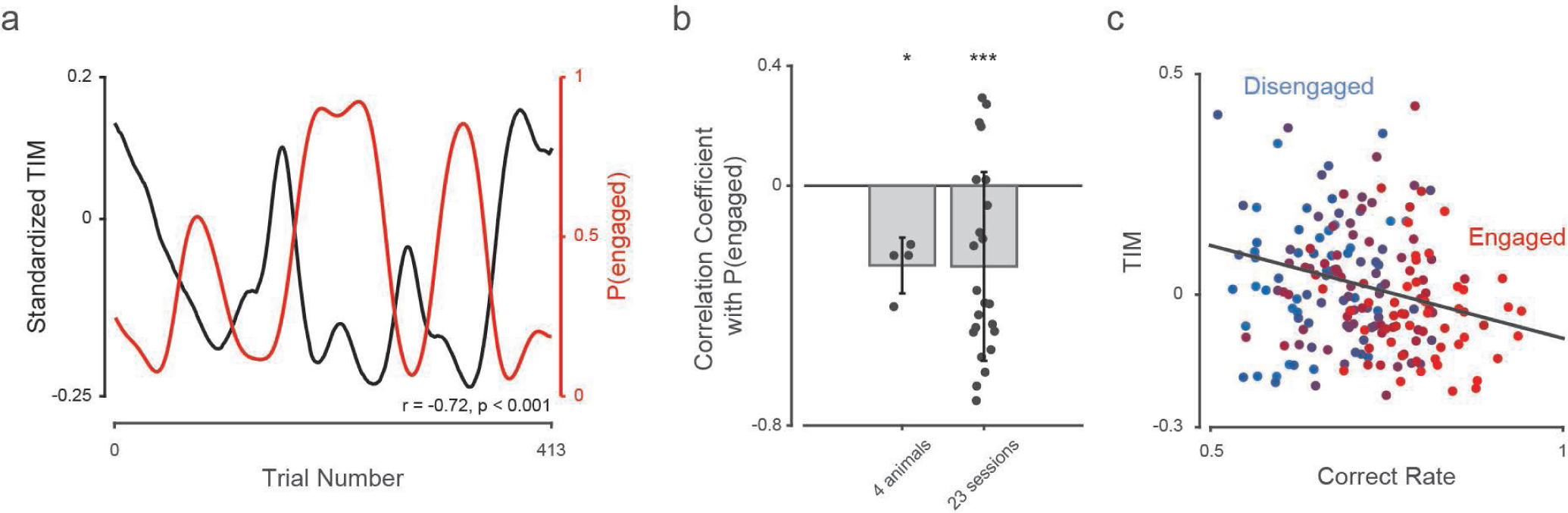
The negative TIM-engagement correlation persists after removing outcome-related regressors from the TIM calculation. **a)** The fluctuation of smoothed TIM and P(engaged) in an example session (same session as in fig. 5c; r = - 0.717, p = 2.114e-66). **b)** The negative TIM-engagement correlation persists at both single-animal and single-session levels (Correlation coefficients of 4 mice = -0.2658 (mean) ± 0.0931 (standard deviation), p of one-sample t-test = 0.0107; Correlation coefficients of 23 sessions = -0.2687 ± 0.3149, p of one-sample t-test = 4.8284e-04). Both TIM and P(engaged) were smoothed with a 50-trial long Gaussian filter before testing the correlation coefficients. **c)** The negative TIM-engagement and TIM-performance correlations persist after removing outcome-related regressors. (TIM-correct rate correlation: r = -0.2520, p = 0.0004, n = 4)

**Supplementary figure 6:**
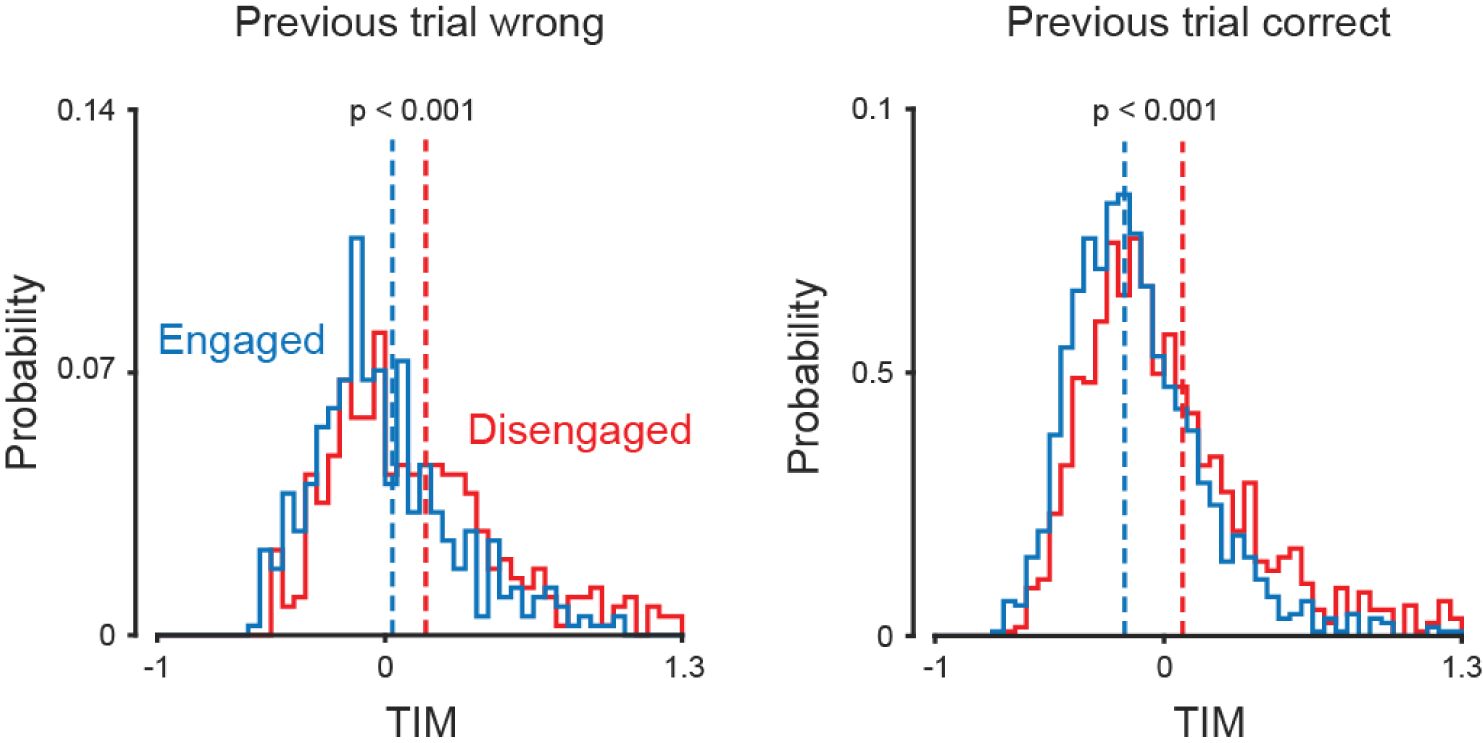
Trials were grouped based on the previous trial’s outcome. Dashed lines denote the mean TIM for engaged and disengaged trials. TIM values were not smoothed across trials. After controlling the outcome of the previous trial, TIM is still higher during disengagement (red dashed line to the right of blue dashed line). (Left: TIM of engaged trials = 0.0317 (mean) ± 0.3965 (standard deviation), TIM of disengaged trials = 0.1783 ± 0.4446, unpaired t-test, p = 1.1484e-06, 794 trials from 4 animals are included; Right: TIM of engaged trials = -0.1108 ± 0.3112, TIM of disengaged trials = 0.0382 ± 0.4219, p of unpaired t-test = 1.5435e-22, 2412 trials from 4 animals are included.)

**Supplementary figure 7:**
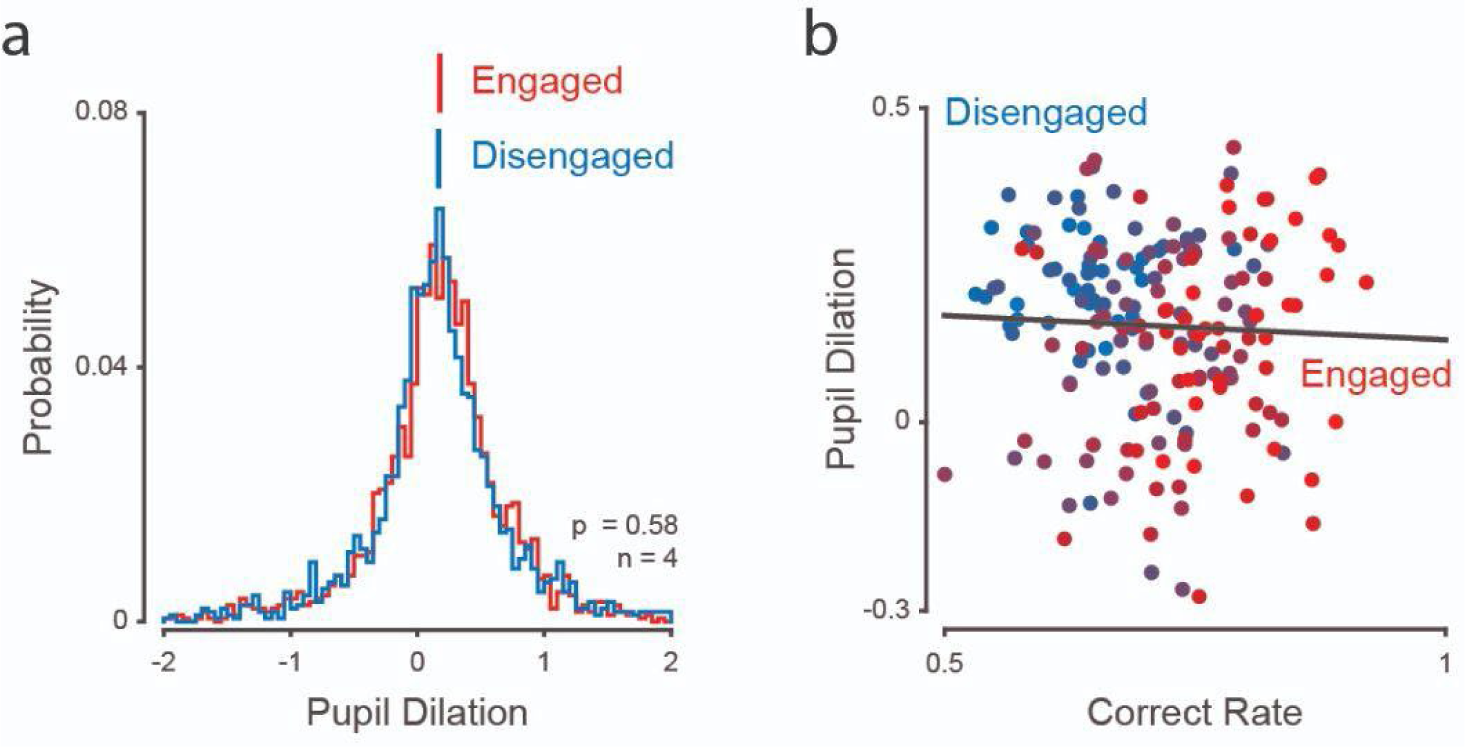
Pupil dilation is not correlated with P(engaged) or task performance. **a)** Distribution of pupil dilation of engaged/disengaged trials. The red/blue lines denote the mean pupil dilation of engaged/disengaged trials respectively. No smoothing across trials was applied. There is no significant difference in pupil dilation (the pupil diameter difference between the stimulus and baseline epochs) between engaged and disengaged states (pupil dilation of engaged trials = 0.1783 (mean) ± 0.5670 (standard deviation), pupil dilation of disengaged trials = 0.1676 ± 0.6217, p of unpaired t-test = 0.5755, n = 4). **b)** Pupil dilation and correct rate were both smoothed with a 50-trial long Gaussian filter and downsampled every 50 trials. Each dot denotes a trial. The color of each dot was assigned based on the value of P(engaged). Pupil dilation does not correlate with task performance (Pupil dilation-correct rate correlation: r = -0.0455, p = 0.5318, n = 4).

**Supplementary figure 8:**
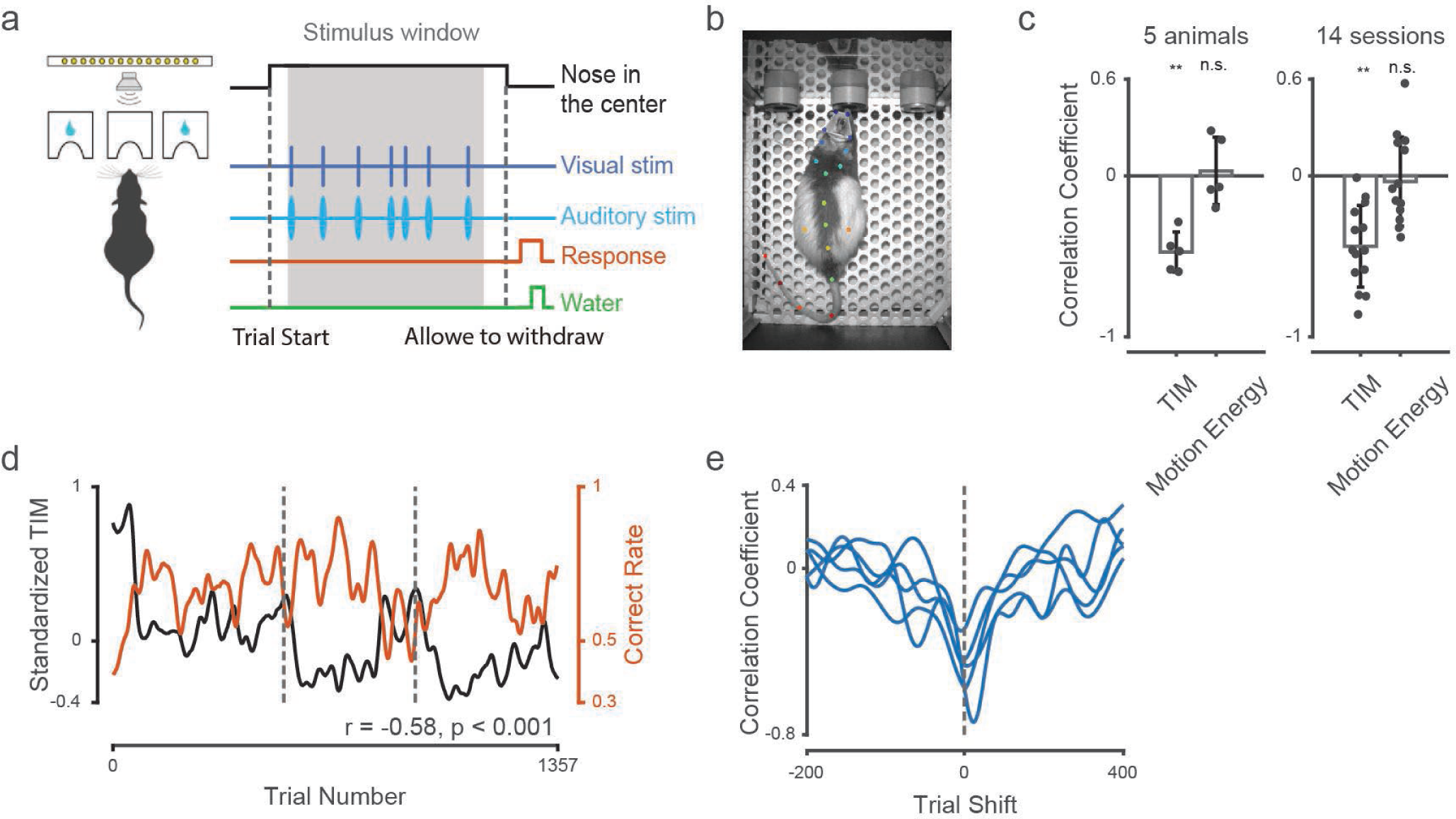
TIM is a negative indicator of performance in a freely-moving rat dataset. **a)** Freely-moving rats were trained to do a multisensory perceptual decision-making task. Animals were trained to report if the number of stimulus events (clicks, flashes, or both) during the stimulus window was higher or lower than a learned category boundary (12 events). Choices to the right (left) port were rewarded when the stimulus was higher (lower) than 12 events. **b)** 19 body parts were tracked with DeepLabCut. Color dots denote DeepLabCut labels. **c)** The correlation between TIM/motion energy and task performance; dots indicate individual animals (left) and sessions (right). Bars show the averaged coefficients across animals and sessions. Both TIM and correct rate were smoothed with a 30-trial long Gaussian filter before calculating the correlation coefficients. We found a negative correlation between TIM and task performance, though the motion energy was independent of task performance. (Left: TIM-correct rate correlation of 5 animals = -0.4734 (mean) ± 0.1254 (standard deviation), p of one-sample t-test= 0.0011; Motion energy-correct rate correlation of 5 animals = 0.0303 ± 0.2098, p of one-sample t-test = 0.7627. Right: TIM-correct rate correlation of 13 sessions = -0.4747 ± 0.0158, p of one-sample t-test = 0.0150; Motion energy-correct rate correlation of 13 sessions = 0.0032 ± 0.2218, p of one-sample t-test = 0.9870) **d)** The fluctuations of TIM and task performance in 3 example sessions from one rat. Both TIM and correct rate were smoothed with a 30-trial long Gaussian filter. (r = -0.5811, p = 2.346e-123) **e)** Trial shift test shows that there is no time lag in the negative TIM-performance correlation. Each line indicates, for a single animal, the correlation coefficient for the time lag indicated on the horizontal axis.

**Supplementary table 1:**
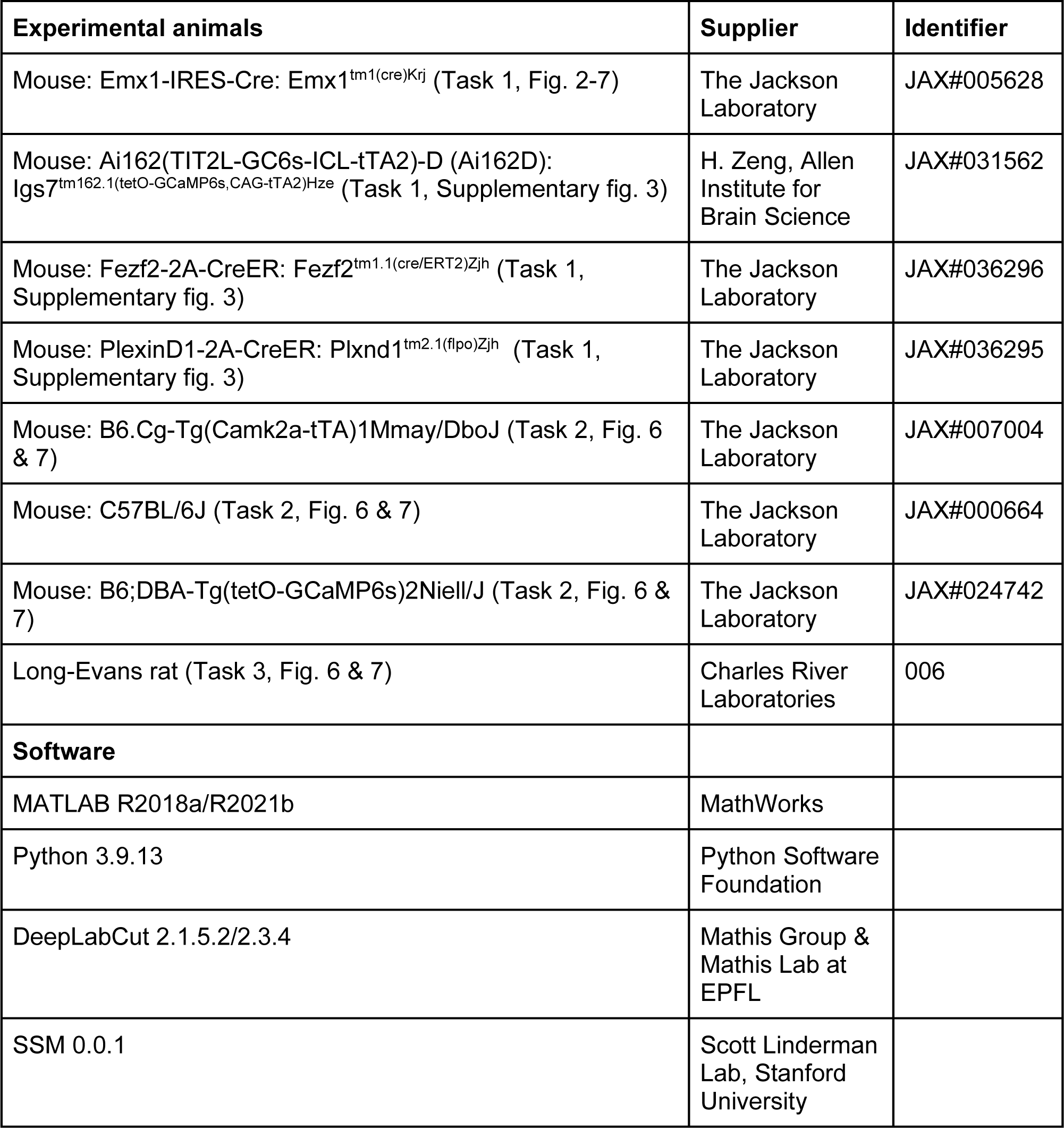
Animal and software information.

## Methods

All surgical and behavioral procedures adhered to the guidelines established by the National Institutes of Health and were approved by the Institutional Animal Care and Use Committee of Cold Spring Harbor Laboratory and the University of California, Los Angeles David Geffen School of Medicine. We used 8-25 week old male and female mice and 23-62 week old male Long-Evans rats. Detailed information on the animal source is provided in Supplementary Table 1. No statistical methods were used to pre-determine sample sizes. Sample sizes are similar to previous publications. Mouse strains were acquired from the Jackson Laboratory, Allen Brain Institute, or generated at Cold Spring Harbor Laboratory. The mouse room had a relative humidity of 30-70%, and room temperature ranging from 69-78°F. To avoid potential aberrant cortical activity patterns, EMX mice were on a doxycycline (DOX)-containing diet, preventing GCaMP6s expression until they were 6 weeks or older.

### General surgical procedures

Surgeries were performed under 1-2% isoflurane in oxygen anesthesia. After induction of anesthesia, 1.2 mg/kg meloxicam was injected subcutaneously and sterile lidocaine ointment was applied topically to the skin incision site. After making a midline cranial incision, the skin was retracted laterally and fixed in position with tissue adhesive (Vetbond, 3M). We then built an outer wall using dental cement (C&B Metabond, Parkell; Ortho-Jet, Lang Dental) along the lateral edge of the dorsal cranium (frontal and parietal bones). A custom titanium skull post was then attached to the dental cement. For skull clearing, the skull was thoroughly cleaned followed by the application of a thin layer of cyanoacrylate (Zap-A-Gap CA+, Pacer technology). Mice were allowed to recover for 5-7 days after the skull clearing before starting the data collection.

### Auditory task for the mice used in widefield imaging (Task 1)

Data used here are from a previously published study^22^. The behavioral setup was controlled with a microcontroller-based (Arduino Due) finite state machine (Bpod r0.5, Sanworks) using custom Matlab code (2015b, Mathworks) running on a Linux PC. Servo motors (Turnigy TGY-306G-HV) and touch sensors were controlled by microcontrollers (Teensy 3.2, PJRC) running custom code. Mice were trained on a delayed, spatial discrimination task. Mice initiated trials by placing their forepaws on at least one of the two handles, which were mounted on servo motors that rotated out of reach during the inter-trial period. To initiate trials, animals placed their forepaws on the handles and, after a variable duration of 0.25-0.75s of continuous contact, the auditory stimulus was presented. Auditory stimuli consisted of a sequence of Poisson-distributed, 3-ms long auditory click sounds, presented from either a left and/or right speaker for a variable duration between 1 and 1.5 s. The stimulus period was followed by a variable delay (up to 1 s), after which servo motors moved two lick spouts close to the animal’s mouth. If the animal licked twice on the side where more sensory events were presented, a drop of water reward was dispensed. After a spout was licked twice, the contralateral spout moved out of reach to force the animal to commit to its decision. The water volume rewarded per trial (typically 1.5 to 3 µL) was constant within a single session but was sometimes adjusted daily based on the animal’s body weight. 5 mice (out of 9) in supplementary figure 4 were used in an optogenetic inactivation experiment based on the same task^22^. To ensure the inactivation always covered the same time window, the durations of stimulus and delay epochs for these mice were fixed to 1.0s and 0.5s respectively. All optogenetic inactivation trials were removed before calculating TIM in case optogenetic manipulation changes the movement pattern.

Data was collected from multiple sensors in the behavioral setup. Touch sensors using a grounding circuit on handles and lick spouts detected contact with the animal’s forepaws and tongue, respectively. A piezo sensor (1740, Adafruit LLC) below the animal’s trunk was used for monitoring body and hindlimb movements. Two webcams (C920 and B920, Logitech) were positioned to capture the animal’s face/upper body (lateral view) and the ventral surface of the body (bottom view). The frame rate of both cameras was 30 Hz. The pupil diameter data in Figure 6a,c was extracted from the lateral videos with customized MATLAB code.

Trained mice were housed in groups of two or more under reverse light cycle (12-hour dark and 12-hour light) and trained during their active dark cycle. Animals were trained over the course of approximately 30-60 days.

### Visual task for the mice used in pupil-engagement analysis (Task 2)

Mice were trained in a visual spatial-temporal discrimination decision making task to monitor pupil dynamics during task engagement. Visual stimuli timing was equivalent to Task 1, described above^22^. Stimuli were presented in a screen (LG LP097QX1, Adafruit) placed 10cm in front of the mouse covering 55 to -55 degrees, calibrated to approximately 50 lux. The eye was recorded through a camera placed at 70 degrees, in the mouse visual field, and 14cm away. The pupil was illuminated with an 850nm LED (IR30, CMVision) and recorded at 30-60Hz using a monochrome camera (CM3-U3-13Y3M-CS, FLIR) equipped with a 12mm lens (NVM-12M23, Navitar) and spacer (ACC-01-5004, FLIR). Custom hardware and software controlled the behavioral task and synchronization with the camera. A micro-controller (Teensy 4.0, PJRC) recorded task events and mouse licking, controlled task timing and emitted a TTL pulse on stimulus presentation that was connected to the general purpose pins of the camera. The status of GPIO was recorded for each frame to recover the precise timing of the stimulus. Data was compressed online using an Nvidia hardware video encoder (through FFMPEG) and stored to disk with custom software (https://github.com/jcouto/labcams).

### Rat behavioral training (Task 3)

Freely moving rats were trained to do a 2-choice auditory/visual decision task based on a task developed previously^61^. Experiments took place in dark, sound-proof rigs (Industrial Acoustics, Bronx, NY). Trials with visual, auditory, and multisensory stimuli were interleaved. In the behavior setup (composed of one speaker, one LED board, and three ports), the rat initiated the trial by putting its nose in the central port. Next, the speaker and/or LED board started to play the stimulus sequence (sound clicks, white light flashes, or both). The number of stimulus events, presented over 1000 ms, ranged from 7 to 17. The decision boundary (12 click/flash stimuli) was abstract and learned with experience by the animal. When the number of stimulus events was above 12, right choices were rewarded; when the number was below 12, left choices were rewarded. The rewarded sides of the trials with exactly 12 stimuli were assigned randomly. The duration of the inter-event interval followed a Poisson distribution. The stimulus window was followed by a short delay period. The duration of the delay period was randomly assigned from an exponential distribution (mean = 0.12 s). After this delay period, an auditory go cue informed the animal that it could leave the central port and make the choice. After the rat poked its nose into the left or right port to report the choice, a reward (24 µL water) or a punishing sound (2.0s long sharp noise) was delivered based on whether the choice was correct. A 2.0 s long white noise sound was delivered if the rat left the central port too early (i.e., during the stimulus or delay period). A webcam (CM3-U3-13Y3M-CS 1/2” Chameleon^®^3 Monochrome Camera) was used to monitor animal movements. The camera was positioned above the animal and its frame rate was 80Hz.

### DeepLabCut tracking and motion energy measurement

To quantify the movement of different body parts, we employed DeepLabCut (version 2.1.5.2 and 2.3.4), a neural-network-based motion tracking software, to track the movement of 27 body parts of the head-fixed mice performing the auditory decision-making task^25^. 14 body parts from the lateral videos and 13 parts from the bottom videos were labeled and tracked (Fig. 4a). We trained two DeepLabCut models (one for the lateral videos, and the other one for the bottom videos) for each of the mice. In each model, about 80-150 frames were extracted and labeled as the training input. The sample frames were selected from different sessions to make the tracking more robust across sessions. After training, we generated labeled videos to test the tracking quality, and kept retraining with new sample frames/refined labels until the tracking quality was visually acceptable (Supplementary videos 1 & 2). To further remove the outliers with poor tracking, we replaced all the values that are > 5 standard deviations away from their mean positions with NaNs.

We next measured the motion energy of DeepLabCut-tracked body parts in the stimulus and delay epochs. Motion energy was defined as the cumulative position change of each body part across frames. Because the lengths of stimulus and delay epochs are variable, the number of trials with the longest stimulus/delay epochs may be small. We omitted all frames that were longer than 70% of the longest stimulus/delay epoch to reduce the noise induced by low trial number. For trials that were shorter than this duration, timepoints after the trial ended were represented with NaNs. We standardized the motion energy of different body parts separately and then averaged all the frames and body parts in each trial together to get a single motion energy value for each trial.

To directly compare the motion energy of engaged and disengaged states, the trials with the highest 20% P(engaged) values of each animal were designated as “engaged trails”. Similarly, the trials with the lowest 20% P(engaged) values were designated as “disengaged trails”. To relate P(engaged) to other behavioral metrics, we first smoothed both P(engaged) with a 50-trial long Gaussian-weighted moving average filter. We then calculated Pearson’s correlation coefficient between the smoothed P(engaged) and the other behavioral metric using the MATLAB function *corrcoef.m*.

### Task Independent Movements (TIM) calculation

Our new metric, TIM, quantifies the magnitude of movements that are independent of task events. This process requires 2 steps. First, we took 7 task variables: current trial’s stimulus strength (*v*_*stim*(*n*)_), choice (*v*_*choice*(*n*)_), outcome (*v*_*reward*(*n*)_), an interaction term of choice and outcome (*v*_*interaction*(*n*)_); the previous trial’s choice (*v*_*choice*(*n*−1)_), outcome (*v*_*reward*(*n*−1)_), and an interaction term of choice and outcome (*v*_*interaction*(*n*−1)_). These 7 variables were used as regressors in a linear model trained to predict the position of each labeled body part (Equation 1). At each video frame (*t*), regressions were performed to predict the x and y values of all body parts (*x^^^*(*t*) or *y^^^*(*t*)). Given the association between engagement and task performance, we balanced the number of correct and incorrect trials before performing the regression to avoid inducing regression quality bias to our analysis. 5-fold cross-validated regression was next performed with MATLAB function *cvpartition.m* and *fitlm.m*. The R^2^ values of regressions were highly variable across mice, indicating that there are significant individual differences in the level of moving stereotypically (Supplementary figure 2).

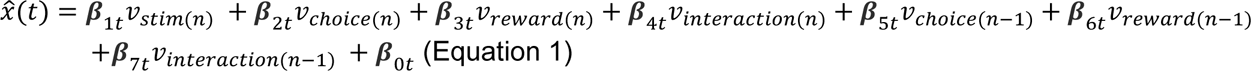

*y^^^*(*t*) was calculated in the same way. Next, we obtained TIM by calculating the Euclidean distance between model-predicted positions and actual positions (Equation 2).

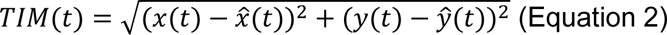

Similar to the motion energy calculation, we restricted calculations of TIM to the stimulus and delay epochs for each body part, and omitted all frames that were longer than 70% of the longest stimulus/delay epoch. The trials with shorter stimulus/delay epochs were filled with NaNs. TIM from each body part was then z-scored to avoid the result being dominated by few body parts. We then averaged TIM across all body parts and frames to get a single value for each trial (Fig. 5b). We also tested the Pearson correlation coefficient of P(engaged) and TIM (Fig. 5c,d). In this case, both P(engaged) and TIM were smoothed with a 50-trial long Gaussian-weighted moving average filter before calculating the correlation coefficients.

### Pupil diameter measurement

The pupil diameter of the mice from which we collected widefield imaging data (figure 6a) was extracted from the lateral videos with customized MATLAB code^23^. We first binarized all the frames and selected the continuous areas bigger than 10 pixels. These areas are considered potential pupil areas. The whiskers sometimes cut the pupil area into multiple pieces in the binarized images. Disconnected areas were merged together to remove this effect. We next used MATLAB function *regionprops.m* to detect the centroids in the merged areas and computed their diameters. Missing values (e.g. due to poor image processing and failed centroid detection) were then extrapolated using forward and reverse autoregressive fits from the remaining samples. Individual pupil diameter traces were then smoothed with a 10-frame long averaging filter. Additionally, data outside six standard deviations were assigned zero weight in smoothing.

For data presented in figure 6c, pupil diameter was extracted using customized software (https://bitbucket.org/jpcouto/mptracker) based on OpenCV^29^. In brief, a region of interest around the pupil was manually selected and used to convert pixels to mm, assuming an eye diameter of 6mm from which ∼80% is visible. The contrast of individual frames was equalized using adaptive histogram equalization. Opening and/or closing morphological operations were done to minimize whisker artifacts and a threshold was applied to the image. Blobs resembling an ellipse with center in the eye were then extracted and the contours fit with an ellipse. The diameter is taken as the diameter of a circle with the same area as the ellipse.

For all the analyses presented in Fig. 6, the pupil was smoothed with a gaussian filter (50 trials long).

### Inferring latent states with the GLM-HMM

To assess the correlation between TIM and states of engagement, we fit a 3-state hidden Markov model with Bernoulli generalized linear model observations (GLM-HMM)^2^. The model is described by a KxK transition matrix (where K represents the number of states) and a set of weights for each state (w_c_^(k)^, where c represents the corresponding Bernouli GLM input parameter and k represents the state). We used two input parameters to the model, stimulus and bias, in order to predict choice behavior. The model was trained using expectation-maximization (EM). Model hyperparameters were first selected with a grid search, where hyperparameter performance was assessed via 10-fold cross validation. Because EM is not guaranteed to converge at a global maximum likelihood, we ran EM 10 times for each fold of cross-validation and picked the model with the highest cross-validated performance. The EM fitting was performed as previously described^2^ using the SSM Python package^62^. We used maximum a posterior (MAP) estimation to estimate the HMM transition matrix and GLM weights. Models were fit to the combined data from all mice because some mice lacked enough data to train individual models. Posterior state probabilities were then inferred via the forward-backward algorithm, also implemented within the SSM package.

### Psychometric function fitting

Psychometric functions were modeled with a four-parameter cumulative Gaussian:

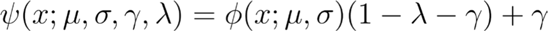

Where *x* describes the stimulus coherence (negative indicates more leftward evidence, positive indicates more rightward evidence), σ describes the inverse slope, *μ* describes the bias, *γ* and *λ* define the low and high lapse rates respectively. *ψ* is the cumulative normal function.

These parameters were estimated using Nelder-Mead Optimization. Psychometric curves were computed from the same trials that were used to train the GLM-HMM. When fitting psychometric curves for the GLM-HMM states, data from all mice were pooled.

### Widefield imaging

Widefield imaging was done as reported previously^22,23,63^. In brief, we used an inverted tandem-lens macroscope and an sCMOS camera (Edge 5.5, PCO). Imaging resolution was 640 x 540 pixels after data were spatially binned at 4x. This resulted in a spatial resolution of about 20 micrometers per pixel. A 525 nm band-pass filter (#86-963, Edmund optics) was used to isolate fluorescence signal. Data were acquired at a framerate of 30 Hz., with alternating blue (470 nm, M470L3, Thorlabs) and violet light (405 nm, M405L3, Thorlabs) delivered along the same excitation path. This enabled isolation of calcium-dependent signals from intrinsic signals, such as hemodynamic responses. The 405 nm violet excitation captures non-calcium dependent GCaMP fluorescence, which can thus be subtracted from the signal obtained with blue (470 nm) excitation. All subsequent analyses were based on this subtracted signal. All frames were rigidly aligned to the Allen CCF (Common Coordinate Framework) using four anatomical landmarks on the surface of the skull.

### Preprocessing of neural data

Neural data were preprocessed as described previously^22^. Briefly, rigid-body registration was used to register frames to the median of the first trial. We then used SVD to compute the top 200 spatial and temporal components of the imaging data. These components comprised at least 95% of the total variance over each recording, and this was done to reduce computational requirements for subsequent analyses.

Imaging data was then aligned to five trial periods: pre-trial, handle grab (trial initiation), stimulus, delay, and outcome. Alignment was required because the duration of trial epochs was randomized to reduce temporal correlation between variables.

### Linear encoding model

The linear encoding model was implemented as previously described^23^. In brief, the linear encoding model included task- and movement-related variables. These variables were assembled into a design matrix containing analog and kernel-based regressors. After assembling the design matrix, the model was fit using ridge regression (with MLE-based estimation of the ridge parameters).

Encoding models were fit to data within single sessions. We fit the encoding model twice for every session: first for trials assigned by the GLM-HMM to the engaged state and then for trials assigned by the GLM-HMM to the disengaged state. Because there were typically more trials in the engaged state, these were randomly downsampled so that the trial numbers in each state for a session were always matched. Sessions with fewer than 50 engaged and 50 disengaged trials were discarded to eliminate models fit to insufficient data.

## Acknowledgements

This work was supported by U19NS123716, a Multi-university Research Grant from the Office of Naval Research (AW911NF-16-1-0368), a grant from the National Eye Institute (R01EY022979) and an NSF-NCS collaborative award (2219946). We thank Anup Khanal for assistance with animal handling and training. We also thank Michael Ryan, Felicia Davatolhagh, Lukas Oesch, Lillian Wilkins, and Letizia Ye for their helpful discussions and comments on the manuscript. We thank Felicia Davatolhagh for performing the citation diversity analysis.

## Diversity statement

Recent work in several fields of science has identified a bias in citation practices such that papers from women and other minority scholars are under-cited relative to the number of such papers in the field. Here we sought to proactively consider choosing references that reflect the diversity of the field in thought, form of contribution, gender, race, ethnicity, and other factors. First, we obtained the predicted gender of the first and last author of each reference by using databases that store the probability of a first name being carried by a woman. By this measure and excluding self-citations to the first and last authors of our current paper), our references contain 6.23% woman(first)/woman(last), 12.77% man/woman, 16.74% woman/man, and 64.25% man/man. This method is limited in that a) names, pronouns, and social media profiles used to construct the databases may not, in every case, be indicative of gender identity and b) it cannot account for intersex, non-binary, or transgender people. Second, we obtained predicted racial/ethnic category of the first and last author of each reference by databases that store the probability of a first and last name being carried by an author of color. By this measure (and excluding self-citations), our references contain 8.79% author of color (first)/author of color(last), 17.82% white author/author of color, 25.37% author of color/white author, and 48.02% white author/white author. This method is limited in that a) names and Florida Voter Data to make the predictions may not be indicative of racial/ethnic identity, and b) it cannot account for Indigenous and mixed-race authors, or those who may face differential biases due to the ambiguous racialization or ethnicization of their names. We look forward to future work that could help us to better understand how to support equitable practices in science.

